# Feedback–feedforward dynamics shape continuous motor learning

**DOI:** 10.1101/2025.04.10.648105

**Authors:** Chen Avraham, Firas Mawase

## Abstract

Many real-world motor skills require continuous control under substantial sensorimotor delays, necessitating the coordination of reactive feedback and predictive feedforward control. How these complementary control processes emerge and interact during the acquisition of a novel continuous motor skill remains unclear. We addressed this question by examining how feedback and feedforward control evolve over five days of continuous visuomotor mirror reversal learning. To directly dissociate feedback- and feedforward-related contributions during learning, we combined frequency-domain analyses of continuous tracking with independent behavioral probes of online feedback responses (brief cursor perturbations) and predictive feedforward control (initial movement direction during point-to-point reaches). We found that learning exhibited both temporal and frequency-dependent shifts in the relative contributions of feedback and feedforward control. Early improvements were dominated by rapid online feedback corrections expressed primarily at low movement frequencies, whereas predictive feedforward control emerged more gradually and became increasingly important at higher frequencies, where delayed feedback is insufficient for accurate control. A recurrent neural network model, in which an adaptive cerebellar-like feedback controller provides a teaching signal to a recurrent cortical predictive controller, reproduced these key behavioral signatures, supporting a mechanistic framework in which feedback-driven error signals facilitate the gradual emergence of predictive control. Together, these findings demonstrate that continuous motor learning arises through distinct but interacting feedback and feedforward processes that co-evolve across practice under the constraints imposed by sensorimotor delays.

## Introduction

Optimal motor performance relies on the integration of feedback and feedforward control processes. Feedback control continuously uses sensory information to detect and correct movement errors online, enabling flexible responses to perturbations and internal variability (Desmurget et al., 1999; Pruszynski et al., 2011; Scott, 2004, 2012; Todorov & Jordan, 2002; Todorov, 2004). However, feedback control is inherently constrained by delays arising from sensory processing and motor execution, limiting its effectiveness during rapid movements (Cameron et al., 2014; Shadmehr et al., 2010). To overcome these delays, the nervous system relies on feedforward control mechanisms that generate predictions about the consequences of motor commands, allowing movements to be planned and executed before sensory feedback becomes available and thereby enabling rapid, smooth motor behavior (Franklin & Wolpert, 2011; Shadmehr et al., 2010; Wolpert & Kawato, 1998; Wolpert & Miall, 1996). Together, these complementary control processes support fast, flexible, and stable motor performance across a wide range of task demands.

While the constraints imposed by sensorimotor delays on feedback and feedforward control are relatively well understood, much less is known about how these complementary control processes develop as a new continuous motor skill is learned. Much of our current understanding of motor learning comes from adaptation paradigms involving discrete point-to-point reaching movements under perturbations such as visuomotor rotations and force fields (Ahmadi-Pajouh et al., 2012; Cunningham, 1989; Morton & Bastian, 2006; Shadmehr & Mussa-Ivaldi, 1994; Wagner & Smith, 2008; Yousif & Diedrichsen, 2012). Although these paradigms have provided important insights into how existing sensorimotor mappings are recalibrated, many natural motor skills require continuous interaction with dynamic environments rather than discrete movements. Activities such as driving, playing a musical instrument, or tracking a moving object require ongoing sensorimotor control in which movement planning and online correction operate simultaneously. How feedback and feedforward control develop and interact during the acquisition of such continuous skills therefore remains an important unresolved question.

Continuous tracking under visuomotor perturbations provides a powerful framework for addressing this question because learning can be characterized simultaneously across a broad range of movement frequencies. Previous work has revealed pronounced frequency-dependent learning during continuous mirror reversal (MR) tracking (Yang et al., 2021), suggesting that the balance between feedback and feedforward control may change over the course of learning. However, how the underlying feedback and feedforward processes themselves evolve across movement frequencies and learning timescales remains unresolved.

Here, we addressed this question by directly dissociating feedback and feedforward control throughout five days of continuous visuomotor MR learning. Participants practiced a continuous MR tracking task while we independently probed feedback control using unexpected cursor perturbations during tracking and feedforward control using interleaved point-to-point reaches without visual cursor feedback. Combining these complementary behavioral measurements with frequency-domain analyses allowed us to characterize how feedback and feedforward control evolved across movement frequencies and learning timescales.

To investigate the architectural mechanisms that could account for these behavioral dynamics, we developed a cortico-cerebellar recurrent neural network (RNN) model in which an adaptive feedback pathway provides teaching signals to a predictive controller (Boven & Cerminara, 2023; Feulner et al., 2025; Pemberton et al., 2024). By integrating complementary behavioral measurements with computational modeling, we tested a mechanistic framework in which adaptive feedback facilitates the gradual emergence of predictive control during continuous motor learning.

## Results

The primary objective of this study was to characterize how feedback and feedforward control evolve during extended practice of a novel continuous motor skill. Participants practiced a continuous MR tracking task over five consecutive days, controlling a cursor to follow a target moving along an unpredictable trajectory (Fig. 1A). The target trajectory was generated by summing seven sinusoids (0.1– 2.15 Hz), with distinct frequency components assigned to the *x*- and *y*-axes. This design enabled tracking performance to be quantified separately across movement frequencies and thereby allowed us to examine how learning depended on movement dynamics.

**Figure 1.**
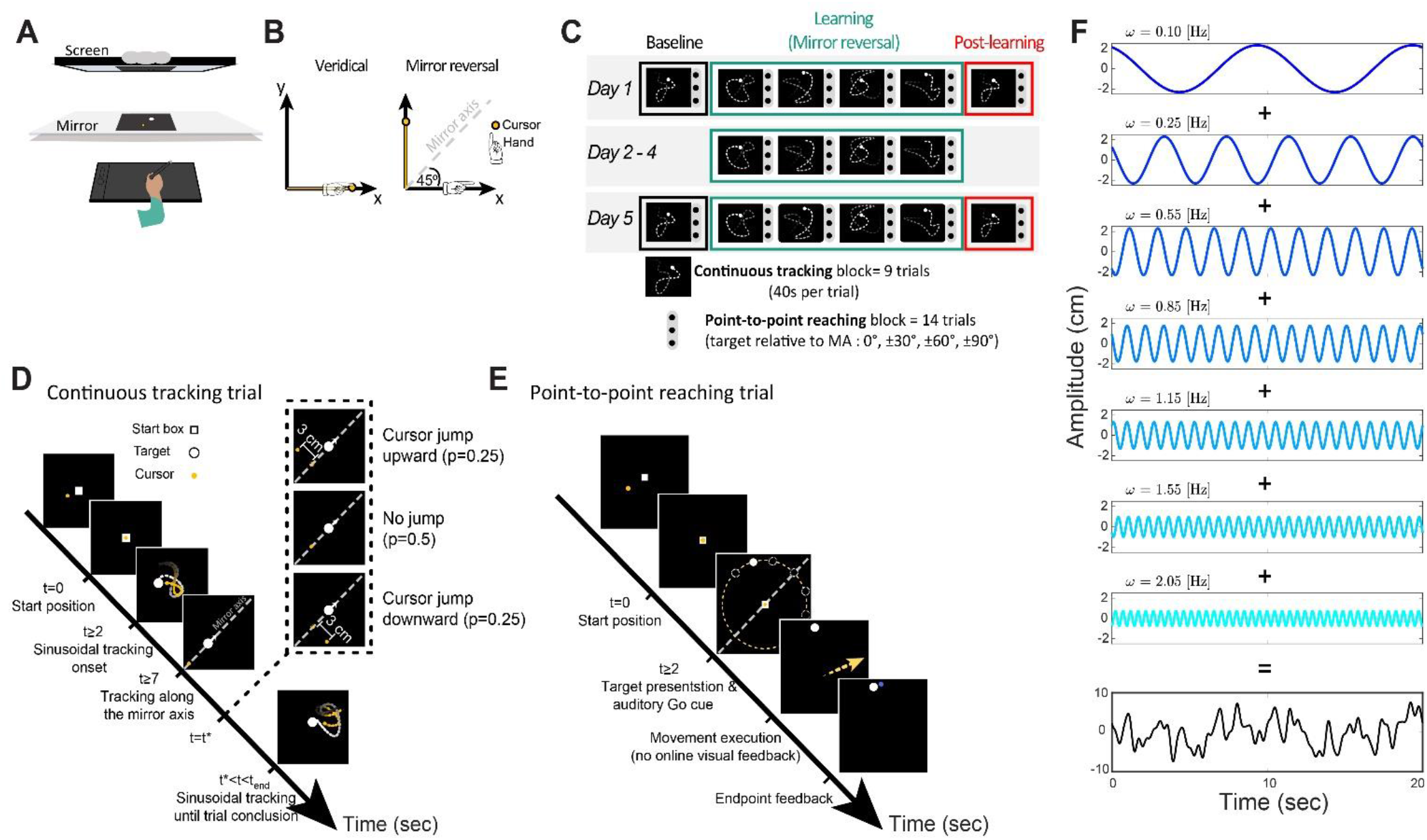
Experimental setup and task design. **(A)** Participants performed a continuous tracking task while seated in front of a horizontally oriented display. The target (white circle) and cursor (yellow circle) were projected onto a semi-reflective mirror, while hand movements were performed with a stylus on a graphic tablet hidden from view beneath the mirror. **(B)** Mapping conditions. Under veridical mapping (left), hand and cursor movements were aligned. Under mirror reversal (MR) mapping (right), movements were reflected across a 45° mirror axis (MA), reversing movement components orthogonal to the MA while preserving movement along the MA itself. **(C)** Experimental timeline. On Days 1 and 5, participants completed baseline (black), MR learning (green), and post-learning (red) continuous tracking blocks. Days 2–4 included only MR learning blocks. Continuous tracking blocks consisted of nine trials. Point-to-point reaching blocks (14 trials each) were interleaved throughout the experiment. **(D)** Continuous tracking trial structure. Each trial began with the cursor positioned inside a start box. After a 2 s hold period, the target initiated sinusoidal motion. At a random time during the trial, the target transitioned to movement along the MA. In half of the trials, an unpredictable vertical cursor jump was introduced, displacing the cursor upward or downward with equal probability. The target then resumed sinusoidal motion until trial completion. **(E)** Point-to-point reaching trial structure. Following a 2 s hold period, a target appeared at one of seven randomly selected locations positioned on a virtual ring (10 cm radius) at angles of 0°, ±30°, ±60°, or ±90° relative to the MA. An auditory Go cue instructed participants to perform a rapid reaching movement. Cursor feedback disappeared once the cursor left the start box to prevent online corrections and reappeared when the movement crossed the target ring, signaling movement termination. The cursor and target remained visible for 2 s before the next trial began. **(F)** Example *x*-axis target trajectory generated by summing seven sinusoidal components with distinct frequencies, amplitudes, and randomized phases, resulting in a complex and unpredictable trajectory.

Participants initially performed the tracking task with veridical cursor feedback to establish baseline performance (Fig. 1B, left). An MR transformation was then introduced around a mirror axis (MA) oriented at 45° (Fig. 1B, right). Under this transformation, hand motion orthogonal to the MA was reversed, whereas motion parallel to the MA remained unchanged. Successfully tracking therefore required participants to learn a new relationship between hand motion and cursor motion.

In addition to characterizing frequency-dependent changes in continuous tracking performance, we used complementary probe trials to independently assess feedback- and feedforward-related control during learning. To probe online feedback control, we intermittently introduced unexpected cursor jumps orthogonal to the MA during tracking and quantified the resulting corrective responses (Fig. 1D; Kasuga et al., 2015; Telgen et al., 2014). To probe predictive feedforward control, we interleaved blocks of rapid point-to-point reaches toward targets presented at different angles relative to the MA, without visual cursor feedback during movement (Fig. 1C). Because the initial movement direction was determined before visual feedback could contribute to online correction, it provided a measure of predictive motor planning under the learned mapping (Fig. 1E).

### Spatial alignment reveals gradual acquisition of the mirror reversal mapping

We first characterized the overall acquisition of the MR mapping by examining the spatial alignment between hand and target movements. A 2 × 2 alignment matrix captured the relationship between hand and target motion along the x- and y-axes (Fig. 2A). Under veridical feedback, perfect tracking corresponds to a unity matrix, reflecting direct correspondence between hand and target motion along each axis. In contrast, the ideal MR matrix exhibits a cross-axis structure reflecting the transformation imposed by the 45° MR.

**Figure 2.**
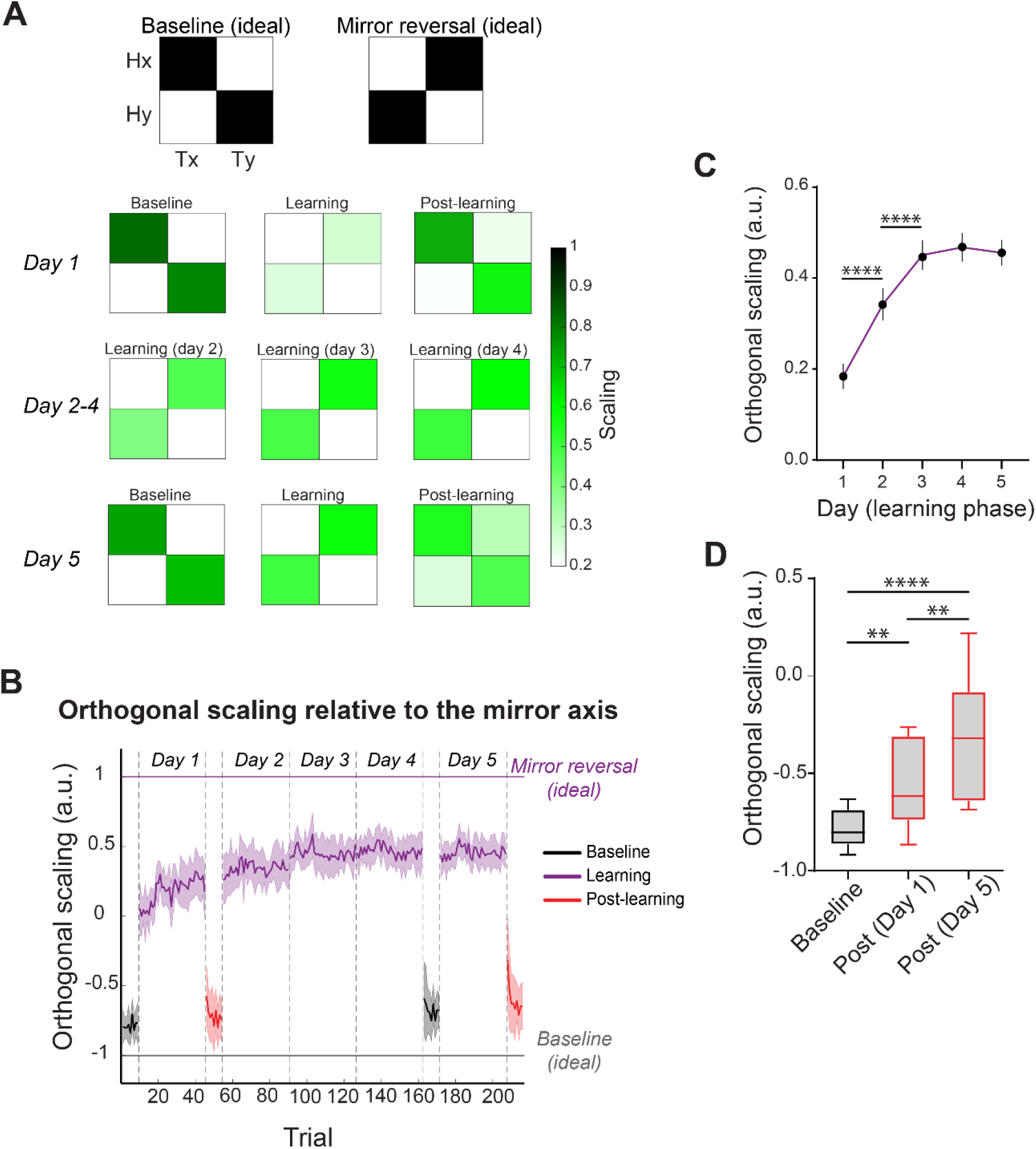
Spatial alignment and orthogonal scaling analysis. **(A)** Ideal alignment matrices representing perfect tracking under veridical (left) and mirror reversal (MR) (right) mappings. Under veridical feedback, the unity matrix reflects a direct correspondence between hand and target movements. Under MR, the ideal matrix reflects the cross-axis transformation imposed by the perturbation. Lower panels show empirically derived alignment matrices averaged across participants. Baseline and learning matrices were additionally averaged across trials within each phase, whereas the Day 1 and Day 5 post-learning matrices represent the first trial following restoration of veridical feedback. **(B)** Trial-by-trial evolution of the orthogonal scaling factor, quantifying the extent to which participants reversed hand movements across the mirror axis (MA). Baseline trials are shown in black, MR learning trials in purple, and post-learning trials in red. A value of −1 reflects the ideal veridical mapping, whereas a value of 1 reflects complete expression of the MR mapping. **(C)** Daily averages of the orthogonal scaling factor during MR learning. The solid line represents the mean across participants and the error bars indicate the SEM. **(D)** Orthogonal scaling factor during baseline (black) and the first post-learning trial on Days 1 and 5 (red), illustrating after-effects following learning. Boxes represent the interquartile range (IQR), with the central line indicating the median. Whiskers extend to the full data range. The corresponding group-averaged alignment matrices for baseline and the first post-learning trial on Days 1 and 5 are shown below. Significance level: ∗∗ *p* < 0.01, ∗∗∗∗ *p* < 0.0001.

During baseline, the empirical alignment matrix closely resembled the veridical unity matrix, indicating accurate tracking under veridical feedback. With practice under the MR mapping, the matrix gradually shifted toward the ideal MR structure. To summarize acquisition of the MR mapping with a single measure, we derived an orthogonal scaling factor from the alignment matrices (see Methods). This measure quantifies the extent to which hand motion is reversed across the MA, with −1 corresponding to the veridical mapping and +1 to complete reversal under the MR mapping. Figure 2B shows the trial-by-trial evolution of the orthogonal scaling factor across practice. At baseline on Day 1, the orthogonal scaling factor was significantly negative (mean ± SEM: −0.79 ± 0.03; one-sample left-tailed *t*-test: *t*(9) = −24.45, *p* < 0.0001), consistent with veridical tracking. The scaling factor increased progressively during practice (Fig. 2D; one-way repeated-measures ANOVA with Greenhouse–Geisser correction: *F*(1.56,14.03) = 69.91, p < 0.0001, 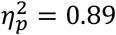). Holm–Šídák-adjusted comparisons revealed significant improvements from Day 1 to Day 2 (*t*(36) = 7.76, *p* < 0.0001) and from Day 2 to Day 3 (*t*(36) = 5.30, *p* < 0.0001). No significant differences were observed beyond Day 3 (Day 3 vs. Day 4: *t*(36) = 0.84, *p* = 0.79; Day 4 vs. Day 5: *t*(36) = 0.58, *p* = 0.81). By the end of practice, the orthogonal scaling factor was significantly positive (final learning block on Day 5: mean ± SEM = 0.46 ± 0.03; one-sample right-tailed *t*-test: *t*(9) = 15.10, p < 0.0001), corresponding to a learning level of 69.17 ± 2.21% (mean ± SEM), demonstrating substantial acquisition of the MR mapping.

To assess persistence of the learned mapping, the MR transformation was removed at the end of learning on Days 1 and 5 and veridical feedback was restored, with participants explicitly informed of the change. Orthogonal scaling differed significantly across baseline and post-learning phases (Fig. 2D; one-way repeated-measures ANOVA: *F*(2, 18) = 21.93, *p* < 0.0001, 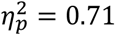). Holm–Šídák-adjusted comparisons revealed significant shifts from baseline following learning on both Day 1 (*t*(18) = 2.98, *p* = 0.008) and Day 5 (*t*(18) = 6.61, *p* < 0.0001), indicating partial persistence of the learned MR mapping after the perturbation was removed. Because orthogonal scaling reflects the overall structure of the alignment matrix, incorporating both diagonal and off-diagonal components, we also quantified after-effects using the mean off-diagonal components alone, as in previous work (Yang et al., 2021). This complementary analysis allowed us to directly compare our results with the conventional off-diagonal measure of remapping. Off-diagonal alignment did not differ significantly between baseline and the first post-learning trial on Day 1 (mean difference = 0.052; Holm–Šídák-adjusted comparison: *t*(18) = 1.95, *p* = 0.067). In contrast, it was significantly greater than baseline on the first post-learning trial of Day 5 (mean difference = 0.147; *t*(18) = 5.54, *p* < 0.001). Thus, although the off-diagonal measure did not detect a significant after-effect after the first day of learning, both analyses indicate that persistence of the learned mapping became more pronounced with extended practice. Notably, orthogonal scaling captured a partial after-effect already on Day 1, suggesting that this measure may be more sensitive to early changes in the learned mapping.

### Continuous mirror reversal learning is frequency dependent

Although the temporal-domain analysis showed clear improvement across practice, it may not fully capture the complexity of the underlying control processes or reveal whether learning was expressed uniformly across the different movement frequencies composing the task. We therefore analyzed performance in the frequency domain using a system-identification approach (Yang et al., 2021). Participants’ tracking behavior was modeled as a linear dynamical system in which sinusoidal target movements elicited sinusoidal hand responses at the same frequencies (Supplementary Fig. S1 and S2). Using discrete Fourier transform, we decomposed hand and target trajectories into their frequency components and constructed two-dimensional gain matrices relating target and hand movements at each frequency pair (see Methods: ‘Frequency analysis’). Figure 3A shows the gain matrices for each frequency, averaged across trials within each experimental phase.

**Figure 3.**
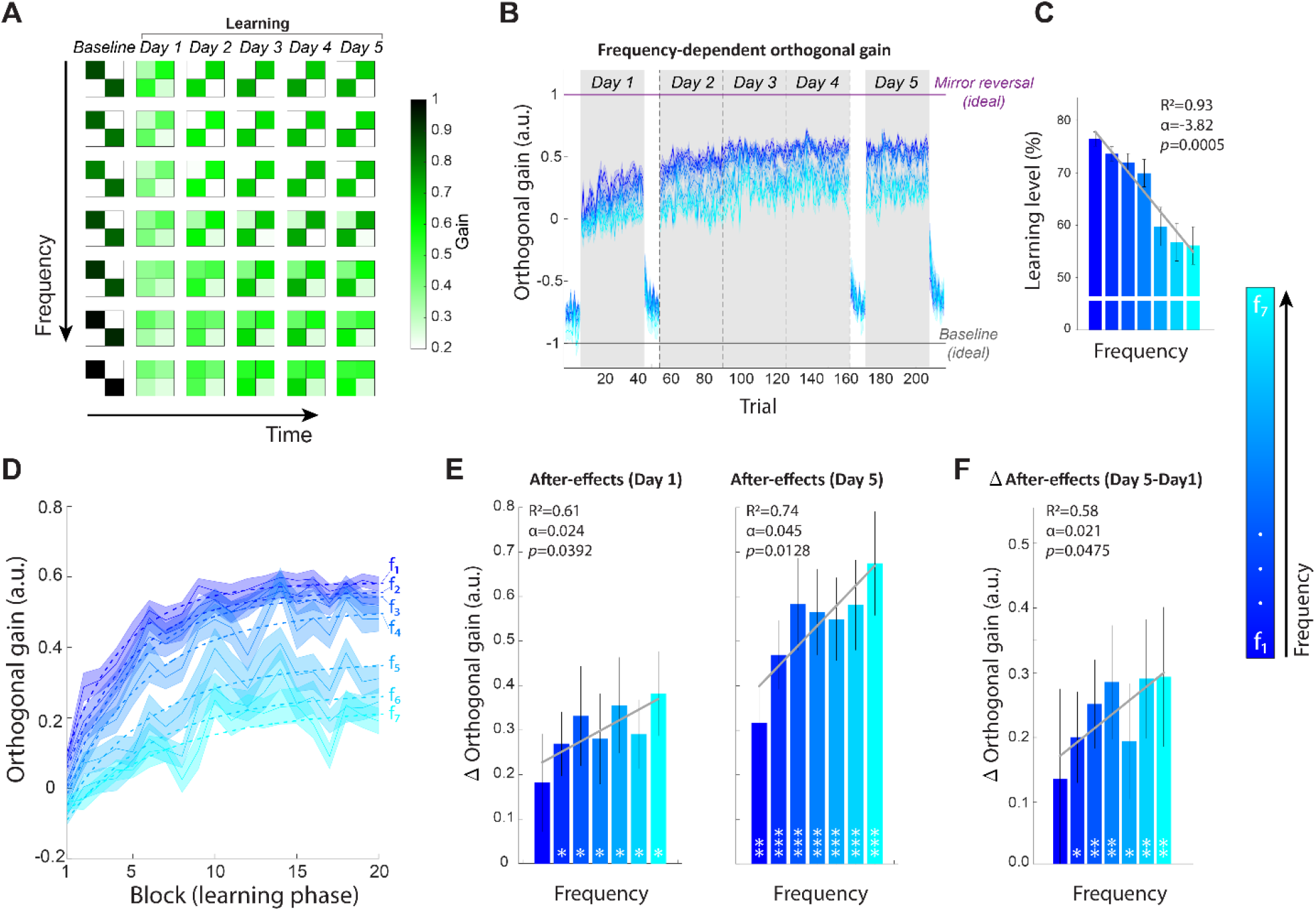
Continuous mirror reversal learning is frequency-dependent. **(A)** Rows correspond to individual frequency components (f_1_-f_7_), ordered from lowest (top) to highest (bottom). The leftmost column shows gain matrices during baseline under veridical mapping on Day 1. Subsequent columns show gain matrices averaged across learning trials for each practice day (Days 1–5) under the mirror reversal (MR) mapping. **(B)** Trial-by-trial changes in orthogonal gain for each movement frequency. Each curve corresponds to a distinct frequency component (f_1_–f_7_). The gray background denotes the learning phase under the MR mapping. An orthogonal gain of −1 reflects ideal performance under the veridical mapping, whereas a value of 1 reflects complete learning of the MR mapping. **(C)** Final learning level as a function of movement frequency. Learning levels were calculated from orthogonal gains averaged across trials in the final learning block on Day 5. Regression analysis revealed a significant inverse relationship between learning level and frequency. **(D)** Learning curves showing block-by-block changes in orthogonal gain during MR learning, plotted separately for each frequency. Dashed lines represent exponential model fits. **(E)** Frequency-specific after-effects on Days 1 (left) and 5 (right), quantified as the difference between orthogonal gain on the first post-learning trial and the corresponding mean baseline gain measured on Day 1. A positive correlation between after-effect magnitude and frequency was observed at both time points. Asterisks denote significant after-effects relative to baseline. **(F)** Changes in after-effects between Day 5 and Day 1. Regression analysis revealed a significant positive relationship between movement frequency and the increase in after-effect magnitude. Asterisks indicate significant increases in after-effects across days. Significance level: ∗ *p* < 0.05, ∗∗ *p* < 0.01, ∗∗∗ *p* < 0.001.

To quantify performance, we calculated the gain orthogonal to the MA from each gain matrix. Analogous to the orthogonal scaling factor derived from the alignment matrices, orthogonal gain quantified how effectively participants inverted their hand movements relative to the MA across the different movement frequencies. Figure 3B displays the trial-by-trial evolution of orthogonal gain averaged across participants, with each curve representing a different target frequency (see Supplementary Fig. S3 for individual participant trajectories). At baseline, orthogonal gains were significantly negative for all frequencies (mean ± SEM, ordered from lowest to highest frequency: −0.79 ± 0.02, −0.75 ± 0.02, −0.72 ± 0.03, −0.75 ± 0.04, −0.75 ± 0.05, −0.79 ± 0.06, −0.89 ± 0.07; Holm–Šídák-adjusted one-sample left-tailed *t*-tests: all *p* < 0.0001), indicating accurate tracking under the veridical mapping across the full frequency range. Despite some variation in baseline orthogonal gain across frequencies, there was no systematic trend across frequency conditions (participant-level linear trend: t(9) = −1.48, *p* = 0.174, *d*_*z*_ = −0.47). The regression across group-averaged values showed a similar pattern (*α* = −0.0145, *F*(1,5) = 2.29, *p* = 0.19, *R*^2^ = 0.31; Supplementary Fig. S4). By the end of practice under the MR perturbation, orthogonal gains increased significantly relative to baseline and became positive (final learning block, mean ± SEM, ordered from lowest to highest frequency: 0.58 ± 0.02, 0.54 ± 0.02, 0.52 ± 0.03, 0.48 ± 0.04, 0.31 ± 0.04, 0.25 ± 0.04, 0.19 ± 0.04; Holm–Šídák-adjusted one-sample right-tailed *t*-tests: all *p* < 0.001), confirming successful reversal of hand movements across the MA at all frequencies. However, the final learning level was frequency dependent (Fig. 3C). Learning levels differed significantly across frequencies (mean ± SEM, ordered from lowest to highest frequency: 76.45 ± 1.35%, 73.63 ± 1.34%, 71.92 ± 1.77%, 69.93 ± 2.61%, 59.74 ± 3.68%, 56.78 ± 3.62%, 56.13 ± 3.59%; one-way repeated-measures ANOVA with Greenhouse–Geisser correction: *F*(1.71,15.42) = 33.80, *p* < 0.001, 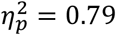), with lower frequencies exhibiting greater learning than higher frequencies. Learning level decreased systematically across the frequency conditions (participant-level linear trend: *t*(9) = −6.82, *p* < 0.001, *d*_*z*_ = −2.16). The regression across group-averaged values showed the same pattern (*α* = −3.816, *F*(1, 5) = 65.93, *p* = 0.0005, *R*^2^ = 0.93), indicating progressively lower learning at higher frequencies. Frequency-domain analysis uncovered distinct learning timescales across frequencies. To quantify learning dynamics, we fitted a single exponential model to the orthogonal gain curves across blocks and extracted the corresponding time constant (*τ*), representing the time required to reach approximately 63% of the learning asymptote. The resulting learning curves and exponential fits are shown in Figure 3D. Time constants increased progressively from low to high frequencies (20.89, 23.82, 24.47, 29.51, 31.84, 32.73, and 38.89 min, respectively), with goodness-of-fit values (*R*^2^) ranging from 0.75 to 0.97, indicating slower learning dynamics at higher movement frequencies.

To determine whether frequency-specific changes acquired during learning persisted after removal of the MR mapping, we quantified after-effects at the end of Days 1 and 5. On Day 1, significant after-effects were observed at all frequencies except for the lowest (Holm–Šídák-adjusted right-tailed one-sample *t*-tests: f2–f_7_: all adjusted *p* ≤ 0.024; f_1_: *t*(9) = 1.65, *p* = 0.067; Fig. 3E, left). By Day 5, significant after-effects were evident across the full frequency range (all adjusted *p* ≤ 0.004; Fig. 3E, right). Interestingly, after-effects were frequency-dependent and increased over time (two-way repeated measures ANOVA: main effect of frequency: *F*(6, 54) = 4.72, *p* = 0.0006, 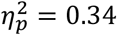; main effect of time: *F*(1, 9) = 11.42, *p* = 0.0081, 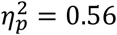; no interaction). Participant-level linear trend analyses showed that after-effects increased significantly across frequency conditions on both Day 1 (*α* = 0.0238 ± 0.0072 SEM, t(9) = 3.29, *p* = 0.0094, *d*_*z*_ = 1.04) and Day 5 (*α* = 0.0452 ± 0.0182 SEM, t(9) = 2.48, *p* = 0.0347, *d*_*z*_ = 0.79). Regressions across group-averaged values showed similar positive relationships on Day 1 (*α* = 0.0238, *F*(1, 5) = 7.69, *p* = 0.0392, *R*^2^ = 0.61) and Day 5 (*α* = 0.0451, *F*(1, 5) = 14.33, *p* = 0.0128, *R*^2^ = 0.74), indicating larger after-effects at higher frequencies.

Post hoc comparisons further showed that after-effects increased significantly from Day 1 to Day 5 at all frequencies except the lowest (Holm-Šídák-adjusted comparisons: f_1_: mean difference = 0.14, *t*(54) = 1.89, *p* = 0.065; f_2_: mean difference = 0.20, *t*(54) = 2.79, *p* = 0.022; f_3_: mean difference = 0.25, *t*(54) = 3.50, *p* = 0.0037; f_4_: mean difference = 0.29, *t*(54) = 3.98 *p* = 0.001; f_5_: mean difference = 0.19, *t*(54) = 2.70, *p* = 0.022; f_6_: mean difference = 0.29, *t*(54) = 4.05, *p* = 0.001; f_7_: mean difference = 0.29, *t*(54) = 4.09, *p* = 0.001) (Fig. 3F). Moreover, at the group-average level, the increase in after-effects across days scaled positively with frequency (linear regression: *α* = 0.0214, *F*(1,5) = 6.83, *p* = 0.0475, *R*^2^ = 0.58), suggesting progressively larger increases over time at higher frequencies.

Together, these results reveal pronounced frequency-dependent differences in the rate, extent, and persistence of MR learning. However, tracking performance alone cannot determine whether these differences arise from distinct changes in feedback and feedforward control. We therefore next examined the independent behavioral probes of these two control processes.

### Feedback control parallels low-frequency learning dynamics

To evaluate the development of feedback control, unexpected cursor jumps were introduced during target tracking (Kasuga et al., 2015; Telgen et al., 2014). These jumps displaced the cursor either upward or downward relative to the MA. Figure 4A illustrates the average corrective hand responses during the 1200 ms following a cursor jump, realigned to reflect an upward displacement. Data are shown for the baseline phase and across learning days (Days 1–5). At baseline, participants produced smooth, well-directed corrective movements that effectively counteracted the displacement, indicating efficient feedback control under familiar conditions. With MR practice, corrective responses gradually shifted toward the pattern expected under the MR transformation, reflecting progressive refinement of feedback control.

**Figure 4.**
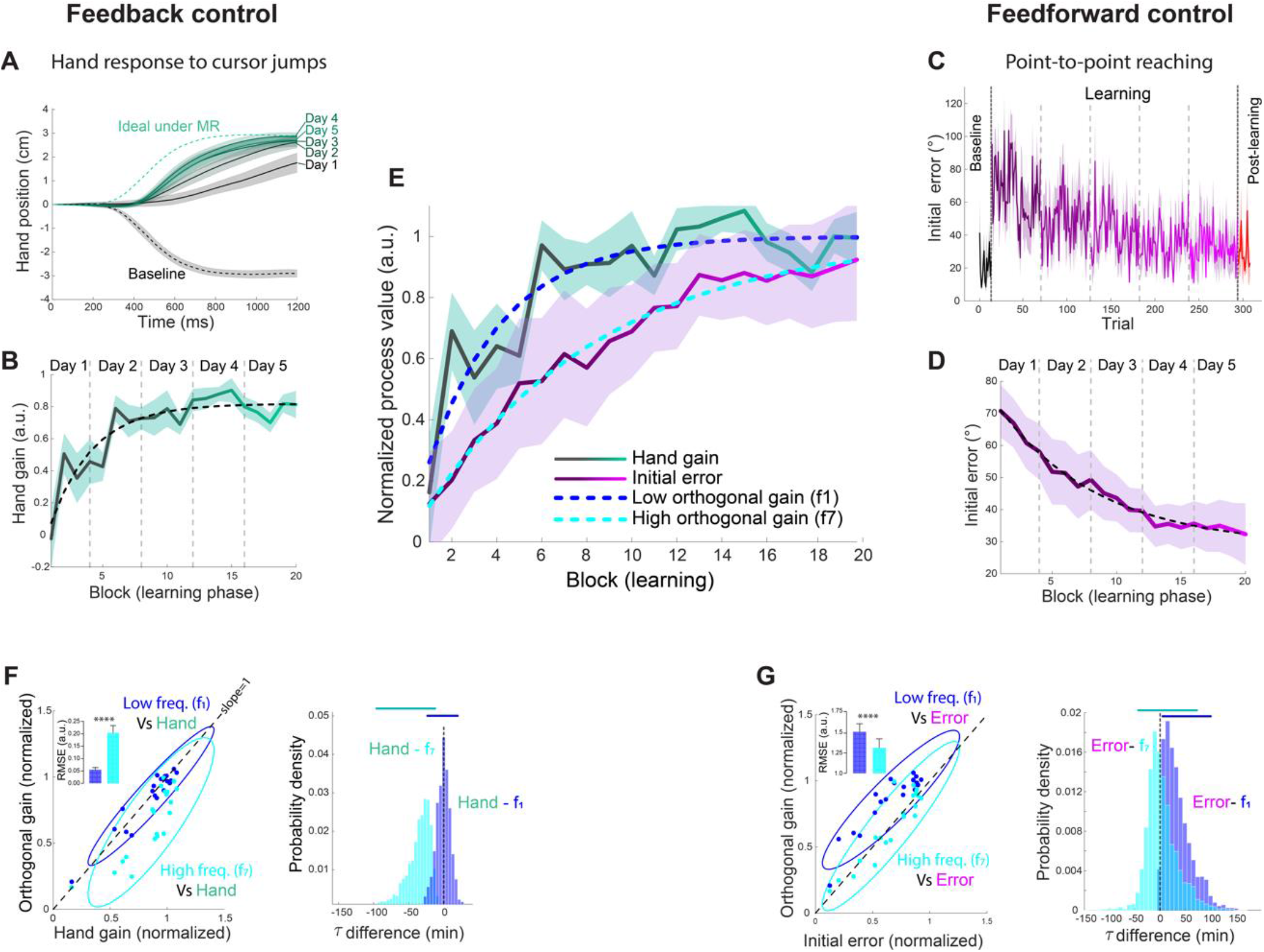
Learning dynamics across feedback- and feedforward-related measures. **(A)** Daily mean hand responses following a 3 cm cursor jump at *t* = 0, aligned to the mirror axis (MA). All responses are displayed as reactions to an upward jump, although both upward and downward displacements were tested. Dashed lines indicate the baseline (black) and ideal mirror reversal (MR) response (green). **(B)** Block-wise changes in hand gain during MR learning. Dashed line indicates the exponential model fit. **(C)** Trial-by-trial initial angular error during point-to-point reaching. Data are shown for the baseline (black), learning (purple), and post-learning (red) phases. **(D)** Block-wise changes in initial angular error during the learning phase, reflecting improvements in feedforward control. Dashed line indicates the exponential model fit. **(E)** Normalized learning curves for hand gain (green) and initial angular error (purple). Dashed lines show normalized exponential fits derived independently from orthogonal gain at low-frequency (f_1_; blue) and high-frequency (f_7_; cyan). **(F)** Left: Relationship between hand gain and orthogonal gain at low-frequency (f_1_; blue) and high-frequency (f_7_; cyan). Each point corresponds to a learning block. The solid black line (y = x) represents perfect correspondence, and ellipses indicate 95% confidence intervals. Inset: RMSE between hand gain and orthogonal gains, shown with SEM error bars. Right: Bootstrapped distributions of time constant (*τ*) differences between the hand gain model and orthogonal gain models. Upper horizontal lines indicate 95% confidence intervals. **(G)** Left: Relationship between initial angular error and orthogonal gain at low-frequency (f_1_; blue) and high-frequency (f_7_; cyan). Each point corresponds to a learning block. The solid black line (y = x) represents perfect correspondence, and ellipses indicate 95% confidence intervals. Inset: RMSE between initial angular error and orthogonal gain models with SEM error bars. Right: Bootstrapped distributions of time constant (*τ*) differences between the initial angular error model and orthogonal gain models. Upper horizontal lines indicate 95% confidence intervals. Across panels A–E, solid lines represent the mean across participants and shaded regions indicate the SEM. Significance level: ∗∗∗∗ *p* < 0.0001.

To quantify these changes, we calculated a hand gain factor for each learning block (Fig. 4B), measuring the degree of alignment with the ideal MR corrective response. A value of –1 indicated baseline alignment, whereas a value of 1 reflected full alignment with the ideal MR response. Across training, hand gain increased significantly, reaching 0.81 ± 0.09 (mean ± SEM) in the final block on Day 5, indicating that corrective responses became strongly aligned with the ideal MR response, despite residual deviations from perfect compensation.

To characterize the temporal dynamics of feedback control, we fitted an exponential model to the averaged hand gain curve (Fig. 4B, dashed line). Figure 4E shows the normalized hand gain alongside orthogonal gains at low and high frequencies (f_1_ and f_7_). The time constant (*τ*) for hand gain was 19.85 min (*R*^2^ = 0.83). Comparison of time constants (Fig. 4F, right) revealed no significant difference between hand gain and low-frequency orthogonal gain (mean difference = −0.51 min; CI = [-19.59, 16.69], *p* = 0.97), indicating similar learning dynamics. In contrast, the time constant for hand gain was significantly smaller than that of the high-frequency orthogonal gain (mean difference = −32.42 min; CI = [-80.82, −10.38], *p* = 0.0013), indicating substantially slower learning at high frequencies. Furthermore, the low-frequency component aligned more closely with the identity line (Fig. 4F, left), as confirmed by a significantly lower root mean squared error (RMSE) relative to the high-frequency component (inset; mean difference = 0.05 ± 0.008 SEM; Two-tailed paired *t*-test: *t*(19) = 5.93, *p* < 0.0001). Together, these findings suggest that the learning dynamics captured by the feedback probe closely parallel those of low-frequency orthogonal gain.

### Feedforward control parallels high-frequency learning dynamics

To assess feedforward control, we tracked the evolution of initial angular error in point-to-point reaching movements that were interleaved between the continuous tracking blocks (Fig. 4C). These short probe trials were designed to allow learning to progress in a largely continuous manner while providing an ongoing readout of predictive control throughout practice. At baseline, the initial error was 21.04 ± 1.18° (mean ± SEM). Upon introduction of the MR, the error increased to 70.73 ± 2.59° in the first block and decreased substantially with practice, reaching 32.33 ± 3.04° in the final MR block, reflecting progressive improvement in predictive control under the MR mapping.

To characterize the dynamics of this process, we fitted an exponential model to the initial error across learning blocks (Fig. 4D, dashed line). Figure 4E shows the normalized initial error alongside orthogonal gains at low and high frequencies (f_1_ and f_7_). The average time constant (*τ*) was 47.19 min (*R*^2^ = 0.98). Comparison of time constants (Fig. 4G, right) revealed no significant difference between initial error and high-frequency orthogonal gain (mean difference = 3.12 min; CI = [–45.11, 76.89], *p* = 0.95). However, initial error exhibited a significantly larger time constant than low-frequency orthogonal gain (mean difference = 34.96 min; CI = [1.01, 101.44], *p* = 0.03), indicating slower development of predictive control relative to low-frequency learning. Figure 4G (left panel) depicts the relationship between initial error and orthogonal gains at low and high frequencies. Here, the high-frequency component aligned more closely with the identity line, as indicated by a significantly lower RMSE relative to the low-frequency component (inset; mean difference = 1.318 ± 0.11 SEM; two-tailed paired *t*-test: *t*(19) = 6.92, *p* < 0.0001). Together, these findings suggest that the learning dynamics captured by the feedforward probe correspond more strongly to those of high-frequency orthogonal gain.

### Cerebellar-like feedback teaches a recurrent cortical controller during continuous mirror reversal learning

To investigate the dynamics between feedback and feedforward processes during continuous motor learning, we developed a recurrent neural network (RNN) model composed of two coupled control pathways: a recurrent cortical-like feedforward controller and a cerebellar-like feedback controller. The recurrent feedforward network generated predictive motor commands based on target kinematics and internal recurrent dynamics, whereas the feedback pathway generated online corrective commands from delayed sensory errors. The feedback loop incorporated a cerebellar-like state estimator that combined delayed visual and proprioceptive information, with delays set to biologically plausible values (*τ*_*vis*_ = 70 *ms, τ*_prop_ = 30 *ms*), to reconstruct the current state of the limb despite sensorimotor delays. Feedforward and feedback commands were summed and passed through simulated arm dynamics to generate cursor motion (Fig. 5A). The model was trained to perform the same continuous two-dimensional MR tracking task used in the behavioral experiment, in which target trajectories were composed of summed sinusoids spanning seven movement frequencies (0.1–2.05 Hz in *x*, 0.15–2.15 Hz in *y*).

**Figure 5.**
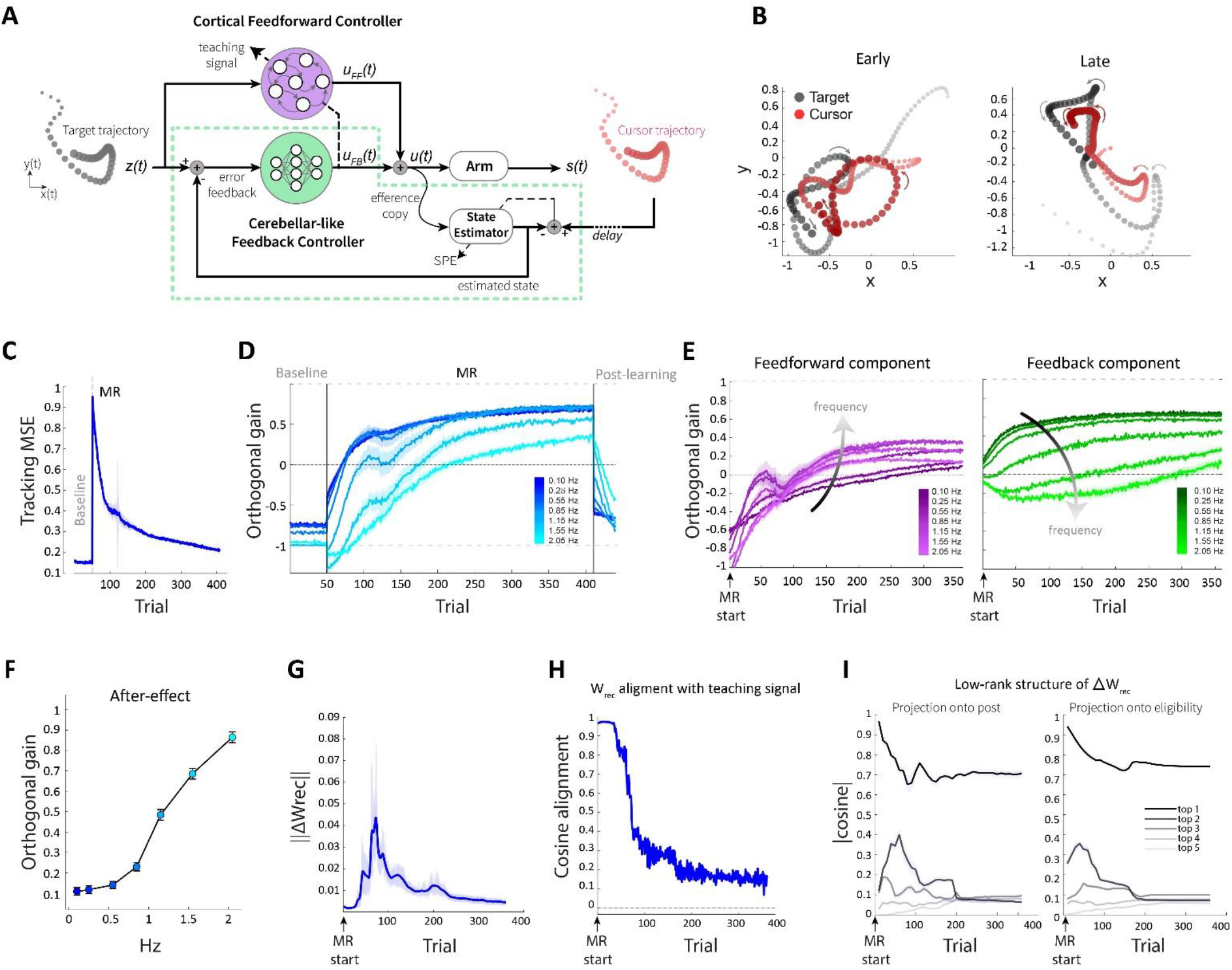
A hybrid feedback–feedforward model reproduces frequency-dependent *dynamics of continuous motor learning*. **(A)** Schematic of the hybrid controller architecture. A cortical recurrent feedforward controller generates a predictive motor command *u*_*FF*_(*t*), while a cerebellar-like feedback controller generates a corrective command *u*_*FB*_(*t*) from sensory error signals. These commands are summed to drive the simulated plant (*u*(*t*) = *u*_*FF*_(*t*) + *u*_*FB*_(*t*)). A state estimator combines efference copy and delayed sensory feedback to compute an estimated state and sensory prediction error (SPE), which supports feedback control and provides a teaching signal to drive cortical plasticity. **(B)** Example target (black) and cursor (red) trajectories during early and late mirror reversal (MR) training. Spatial units are expressed in decimeters (dm; 1 dm = 10 cm). **(C)** Tracking mean squared error (MSE; dm^2^) between target and cursor trajectories across trials. **(D)** Trial-by-trial changes in orthogonal gain plotted separately for each movement frequency. **(E)** Decomposition of orthogonal gain into feedforward and feedback pathway contributions across learning, shown separately for each movement frequency. Left: feedforward (cortical-like) component. Right: feedback (cerebellar-like) component. **(F)** Frequency-dependent after-effects. **(G)** Magnitude of cortical recurrent plasticity across learning, quantified by the norm of the recurrent weight update. **(H)** Alignment between recurrent plasticity and the feedback-derived teaching signal, quantified by cosine similarity. **(I)** Low-rank structure of recurrent weight changes. Projections of the leading modes of *ΔW*_*rec*_ onto postsynaptic (teaching-related; left) and presynaptic (eligibility-related; right) directions.

During pre-training and baseline trials under veridical feedback, the model learned to track the target accurately, achieving stable low tracking error (MSE ≈ 0.15 dm^2^), indicative of effective online control (Fig. 5B&C). Introduction of the MR perturbation produced an immediate disruption in performance, with cursor trajectories deviating strongly from the target (Fig. 5B, left) and tracking error increasing sharply (MSE ≈ 0.95 *dm*^2^). Tracking error then decreased rapidly over the first tens of MR trials, followed by more gradual improvement across later training, ultimately approaching near-baseline levels by the end of practice (MSE ≈ 0.2 dm^2^; Fig. 5C). Consistent with this recovery, cursor trajectories progressively overlapped the target path (Fig. 5B, right).

Next, we examined how tracking performance evolved across movement frequencies. As in the behavioral experiment, orthogonal gain was computed separately for each target frequency component. During MR practice, orthogonal gains progressively shifted from negative values associated with baseline tracking toward positive values consistent with successful MR learning, but this transition exhibited a pronounced frequency dependence: lower frequencies improved more rapidly and reached substantially higher learning levels than higher frequencies. By mid-training, the lowest-frequency components approached orthogonal-gain values of approximately 0.68, whereas higher-frequency components exhibited more limited improvement, reaching values of approximately 0.34–0.56 (Fig. 5D). Critically, frequency-dependent structure was also evident during the post-learning phase. When veridical feedback was restored, low-frequency components rapidly returned toward baseline, whereas high-frequency components exhibited larger and more persistent after-effects (Fig. 5D and 5F). This dissociation indicates stronger persistence of the learned policy at higher frequencies, consistent with the behavioral findings.

To directly examine how the two control pathways contributed across learning, we decomposed orthogonal gain into feedforward- and feedback-specific components (Fig. 5E). This analysis revealed a clear division of labor between pathways: Early improvements were dominated by the feedback pathway, whose contribution increased rapidly and was expressed primarily at low frequencies (Fig. 5E, right). In contrast, the feedforward pathway initially contributed minimally but strengthened progressively across practice, with its contribution becoming increasingly prominent at higher frequencies, where sensorimotor delays fundamentally limit online feedback control (Fig. 5E, left panel). This temporal and frequency-dependent transition closely reproduced the dissociation observed behaviorally in human participants, supporting the conclusion that feedback and feedforward processes contribute differently across learning and movement frequencies. Learning-related improvements were accompanied by structured changes in the recurrent cortical network (Fig. 5G–I). The magnitude of recurrent plasticity increased sharply following introduction of the MR and then gradually decreased as performance stabilized, indicating that the largest recurrent weight updates occurred early during practice (Fig. 5G). To determine whether these recurrent changes were shaped by the feedback-derived teaching signal, we quantified the alignment between recurrent plasticity updates and the teaching-rule update. Recurrent weight changes showed strong initial alignment with the teaching signal, which progressively decreased across practice (Fig. 5H), suggesting that feedback-driven instructive signals strongly shaped early cortical plasticity, whereas later updates became smaller and less consistently aligned with the teaching direction. Analysis of recurrent weight structure further revealed that learning-induced changes were strongly concentrated within a low-dimensional structure (Fig. 5I). The leading one to two modes of recurrent plasticity accounted for most of the projection onto both postsynaptic teaching-related directions and presynaptic eligibility-related directions, whereas higher-order modes contributed minimally. Thus, recurrent reorganization was dominated by a small number of coordinated patterns of synaptic change rather than diffuse modifications distributed across the full recurrent network. This low-dimensional structure suggests that feedback-driven teaching signals constrain cortical plasticity along specific directions, conceptually consistent with reports that learning-related neural activity evolves within low-dimensional manifolds in motor cortex. These findings suggest that the cerebellar-like feedback pathway not only provides online corrections but also supplies structured instructive signals that shape the gradual emergence of predictive recurrent dynamics.

To identify the model components required to reproduce the human-like learning trajectory, we performed three ablation probes. First, removing the feedback-derived teaching pathway while freezing recurrent cortical plasticity preserved limited online compensation through feedback, but strongly attenuated the emergence of the predictive component and substantially reduced high-frequency after-effects (Fig. S5A), indicating that feedback-driven recurrent learning is required for the gradual development of predictive control. Second, replacing the local feedback-driven recurrent learning rule with backpropagation-based optimization improved overall task performance, but failed to reproduce the characteristic frequency-dependent progression and persistence observed behaviorally (Fig. S5B). Thus, generic error minimization alone was insufficient to capture the human-like learning dynamics, supporting the idea that the tutor-like feedback-driven learning mechanism contributes critically to the observed temporal and frequency-dependent structure of learning. Third, ablating the state estimator produced a distinct qualitative impairment in the feedback pathway. During MR learning, the feedback component became strongly negative at the highest frequencies, indicating that delayed sensory errors can drive corrective commands in an inappropriate direction when they are not supported by predictive state estimation (Fig. S5C). This manipulation also exaggerated high-frequency after-effects, suggesting that predictive state estimation contributes to stabilizing learning in the high-frequency regime.

Together, these ablation probes support a model in which feedback serves dual roles during continuous motor learning: stabilizing behavior online through closed-loop correction while simultaneously providing instructive signals that guide the gradual emergence of predictive recurrent dynamics. In parallel, predictive state estimation is required to maintain stable feedback control under sensory delay, particularly at high movement frequencies.

## Discussion

The present study examined how feedback- and feedforward-related control processes evolve during extended practice of a continuous visuomotor MR task. Across behavioral measures and computational modeling, learning exhibited a consistent temporal and frequency-dependent organization. Performance improved more rapidly and reached higher levels at lower movement frequencies, where learning closely paralleled the rapid evolution of feedback-related control. In contrast, higher-frequency learning developed more slowly and more closely paralleled the gradual emergence of predictive feedforward control. Together, these findings suggest that extended continuous skill acquisition is supported by learning processes operating over multiple timescales, with the relative contributions of feedback and feedforward control evolving over practice.

This frequency-dependent organization may reflect changes in the utility of delayed sensory feedback across movement timescales. During slower movements, delayed sensory information can be processed in time to support effective online error correction. As movement dynamics become faster, however, sensorimotor delays increasingly limit the capacity of sensory feedback to modify the ongoing movement, thereby reducing the effectiveness of reactive control and increasing reliance on predictive mechanisms (McNamee & Wolpert, 2019). Previous accounts of how feedback and predictive control change during learning have been developed largely from motor-adaptation paradigms involving discrete, point-to-point reaches (Mathew & Crevecoeur, 2021; Shadmehr et al., 2010). The present findings extend this work to the prolonged acquisition of a continuous skill by showing that movement frequency systematically shapes both the time course of learning and the relative expression of feedback- and feedforward-related control across practice.

Beyond their frequency-dependent relationship, feedback- and feedforward-related measures exhibited distinct learning trajectories. Corrective responses to cursor perturbations changed rapidly during early practice, suggesting that feedback control was quickly recalibrated to the novel sensorimotor mapping. In contrast, feedforward-related changes in the initial direction of point-to-point reaches emerged more gradually, suggesting that the development of an appropriate predictive control policy required more extensive experience. This temporal ordering is compatible with evidence from discrete force-field adaptation paradigms showing that errors initially evoke feedback-based corrective responses and that components of these responses are progressively incorporated into anticipatory motor commands (Albert & Shadmehr, 2016; Milner & Franklin, 2005; Thoroughman & Shadmehr, 1999). Related work also shows that feedback gains adapt to anticipated task dynamics and that feedback responses can change during the early stages of adaptation (Ahmadi-Pajouh et al., 2012; Coltman & Gribble, 2020). The present findings extend this literature by demonstrating distinct feedback- and feedforward-related learning trajectories across prolonged acquisition of a continuous MR skill rather than short-term adaptation of discrete reaches.

An important finding of the present study was the persistence of MR-related behavior after veridical feedback was restored. Orthogonal scaling, which captures the overall expression of the learned reversal, revealed significant after-effects following both the first and fifth days of learning. In contrast, the off-diagonal components of the alignment matrix, the measure used by Yang et al. (2021) to quantify MR cross-axis coupling, provided little evidence of an after-effect after Day 1 but showed a robust effect after Day 5. Thus, convergence between the two measures following extended practice provided stronger evidence that post-learning behavior retained the cross-axis structure of the MR mapping.

MR learning has commonly been characterized as *de novo* learning, in which successful compensation depends primarily on acquiring a new, task-specific control policy rather than recalibrating an existing sensorimotor mapping (Telgen et al., 2014; Yang et al., 2021). This interpretation has been supported in part by the small or absent after-effects observed following removal of the transformation. The robust after-effects observed here following extended practice do not necessarily challenge the proposal that MR compensation was achieved primarily through acquisition of a new control policy. Rather, they show that the learned MR mapping continued to influence behavior after veridical feedback was restored. This persistence could arise from residual or generalized expression of the learned MR policy, from adaptation-like modifications of pre-existing sensorimotor mappings, or from a combination of these processes. Previous work also indicates that conventional error-driven adaptation can be engaged during MR learning even when its effects are inappropriate for compensating the mirror transformation, suggesting that adaptation-related changes may coexist with acquisition of an MR-appropriate policy (Hadjiosif et al., 2021; Wang & Taylor, 2021). The present behavioral measures do not distinguish among these possibilities. Importantly, the after-effects measured here differ from their typical assessment in discrete adaptation paradigms. Classic adaptation after-effects are commonly assessed from the initial direction of movements performed under conditions that minimize visual feedback, providing a relatively direct assay of persistent feedforward recalibration. In contrast, here after-effects were quantified during continuous tracking with veridical visual feedback restored. They therefore reflect the net expression of prior MR learning within a closed-loop control setting, in which residual MR-related output can be corrected online and behavior can progressively return toward the veridical mapping. Accordingly, these after-effects should not be interpreted as a pure measure of implicit adaptation or as direct evidence that the baseline controller was recalibrated.

Although after-effects alone cannot identify the underlying learning mechanism, their frequency-dependent structure is informative when considered alongside the independent feedback and feedforward probes. After-effects were relatively small at the lowest movement frequencies and increased with frequency, with more robust persistence evident following extended practice. One possible explanation is that, after restoration of veridical feedback, online corrective control can more effectively reduce the behavioral expression of residual MR-related output at lower frequencies. At higher frequencies, where sensorimotor delays place greater constraints on online correction, residual expression of the learned transformation may be less effectively counteracted. Notably, this frequency-dependent pattern parallels the control processes measured independently during learning: low-frequency performance more closely followed the rapid evolution of perturbation-evoked feedback responses, whereas higher-frequency performance more closely followed the gradual development of predictive initial movement directions. Thus, the larger after-effects at higher frequencies are consistent with greater persistence of predictive components of the learned control policy, while also reflecting the reduced capacity of restored veridical feedback to counteract their behavioral expression.

The temporal ordering and frequency-dependent relationship between the feedback- and feedforward-related changes raise the possibility that these processes might interact during learning rather than developing entirely independently. More specifically, the rapid emergence of feedback-related changes, followed by the slower development of predictive behavior and high-frequency learning, is consistent with a feedback-scaffold account in which early feedback-based corrections facilitate the gradual emergence of robust predictive control during continuous motor learning. This interpretation is consistent with evidence from sensorimotor adaptation studies that feedback-related muscle responses are progressively modified during learning and that the resulting anticipatory commands share temporal and amplitude-related features with these earlier corrective responses (Albert & Shadmehr, 2016; Maeda et al., 2020; Milner & Franklin, 2005; Thoroughman & Shadmehr, 1999). Furthermore, Feulner et al. (2025) provided computational support for this account by showing that a recurrent neural network trained with an error-based feedback signal developed feedforward-like dynamics over time, mirroring changes in neural population activity in monkey primary motor cortex during motor adaptation.

The computational model presented in this work provided a mechanistic test of this feedback-scaffold mechanism. Within its closed-loop architecture, an adaptive cerebellar-like feedback controller stabilized behavior in the presence of sensory delays while supplying error-related teaching signals to a recurrent cortical-like predictive controller. These signals progressively shaped predictive dynamics that compensated for the MR perturbation, demonstrating how rapid online correction could contribute to the slower acquisition of predictive control. This architecture reproduced key temporal and frequency-dependent features of MR learning observed behaviorally, including the rapid improvement at lower frequencies and the slower development of higher-frequency performance. Disrupting the teaching pathway substantially impaired learning and prevented the model from reproducing this characteristic frequency-dependent progression. This result identifies feedback-derived teaching signals as a critical component of the model’s gradual development of predictive control and provides a computational proof of principle for the proposed feedback-scaffold mechanism. This interpretation is consistent with computational accounts proposing that cerebellar output supports rapid task acquisition and shapes the subsequent consolidation of task-related information in cortical circuitry (Boven & Cerminara, 2023; Pemberton et al., 2024). More broadly, neurophysiological evidence indicates that cerebellar–motor-cortical contributions change across skill acquisition, with cerebellar-related physiological changes evident early in learning and M1 plasticity-like changes emerging after more extensive practice (Spampinato & Celnik, 2017).

Analysis of the recurrent network provided further insight into how this learning was implemented in the model. During the early stages of learning, recurrent plasticity was strongly aligned with the feedback-derived teaching signal, but this alignment weakened as performance stabilized. At the same time, recurrent reorganization remained concentrated within a low-dimensional subspace, indicating that learning was driven by a small number of coordinated modifications rather than diffuse changes throughout the network. This organization is compatible with dynamical-systems accounts proposing that motor learning relies on a limited set of coordinated changes constrained by existing neural population dynamics (Gurnani et al., 2026).

Importantly, the present experimental design was optimized to characterize the temporal and frequency-dependent evolution of feedback- and feedforward-related control, rather than to establish a causal dependency between them. The behavioral data revealed a consistent temporal ordering and correspondence with learning at different movement frequencies, but they do not determine whether feedback-related changes instruct subsequent predictive learning or whether both processes adapt in parallel to shared error signals. The computational model demonstrates that a feedback-scaffold mechanism is sufficient to reproduce the observed behavioral organization, but it does not exclude alternative architectures in which feedback and predictive control develop independently or interact through a different mechanism. Distinguishing among these possibilities will require experiments that selectively manipulate these processes during continuous motor learning.

In summary, the present study shows that continuous MR learning emerges through distinct but interacting feedback and feedforward control processes, and that their relative contributions vary across practice and movement frequencies. By combining continuous tracking, perturbation-based behavioral probes, and computational modeling, we found that lower-frequency learning developed rapidly and closely paralleled changes in feedback-related corrective responses, whereas higher-frequency learning emerged more gradually and more closely tracked the development of predictive control. This organization is consistent with the decreasing utility of delayed sensory feedback at faster movement timescales. A recurrent neural network model with interacting cerebellar-like feedback and cortical-like predictive components captured these dynamics and provided a computational proof of principle for a feedback-scaffold framework in which adaptive feedback both stabilizes behavior and shapes the gradual emergence of predictive control. Together, these findings provide a framework for understanding how reactive and predictive control processes co-evolve during extended acquisition of a continuous motor skill.

## Methods

### Participants

Ten healthy adults (6 female, mean age 26.6 ± 4.32 years) participated in the study. All participants reported no history of neurological or cognitive impairments. Handedness was assessed using the 10-item Edinburgh Handedness Inventory, indicating a strong right-handed preference across participants (mean score: 90.56 ± 16.27). All participants completed the full longitudinal protocol and received monetary compensation (250 ₪) upon completion of the study. Written informed consent was obtained from all participants prior to participation, and the study protocol was approved by the Technion Institutional Review Board.

### Manual interface and data acquisition

The experimental setup consisted of a fixed chair positioned in front of a horizontally oriented computer screen (48 cm width; 1280 × 1024 pixels; 144 Hz refresh rate). Visual stimuli presented on the screen were reflected onto a horizontal semi-reflective mirror placed above the participant’s hand (Fig. 1A). Participants rested their heads against a head support and viewed the stimuli exclusively through the mirror. An opaque fabric placed beneath the mirror prevented direct observation of the hand and forearm, ensuring that participants relied solely on the reflected visual stimuli. Participants performed the task using their dominant hand by holding a stylus pen and moving it across a horizontal graphic tablet (62 × 46 cm; Intuos4, Wacom Co., Japan). The task was implemented using custom MATLAB scripts (The MathWorks Inc., Natick, MA, USA), and kinematic data were sampled at approximately 100 Hz for offline analysis.

### Block design

The longitudinal experiment spanned five consecutive days and included continuous tracking blocks, each comprising nine trials (Fig. 1C). In each trial, participants tracked a target moving along a pseudo-random sum-of-sines trajectory composed of multiple frequency components (Fig. 1F). On Day 1, participants received written instructions and familiarized themselves with the task interface. They then completed a baseline phase consisting of a single tracking block with veridical feedback to establish a reference level of motor performance. Following baseline, an MR perturbation was applied to the cursor, with the MA oriented at 45° relative to the positive *x*-axis (Fig. 1B). Under this condition, participants completed four tracking blocks. At the end of Day 1, the MR perturbation was removed, and participants completed a single post-learning tracking block with veridical feedback to assess after-effects. Days 2–4 focused exclusively on MR learning, during which participants completed four tracking blocks per day. Day 5 replicated the structure of Day 1, including a baseline assessment, MR learning blocks, and a final post-learning block. Participants completed approximately 24 min of MR tracking each day, resulting in a total of approximately 120 min across the experiment. Participants were informed when the MR perturbation was introduced and removed, but were not explicitly informed about the nature of the perturbation itself. To minimize potential temporal effects on performance, training sessions were conducted at approximately the same time each day.

In addition to continuous tracking, participants completed point-to-point reaching blocks interleaved between tracking blocks (Fig. 1C). Within each block, participants performed 14 reaching trials toward targets presented at multiple directions relative to the MA. The initial movement direction in these trials was used to assess the development of predictive feedforward control over the course of learning.

### Continuous tracking trial

Each trial began with a white square start box (10 mm) presented at the center of a black workspace (21.6 × 21.6 cm; Fig. 1D). A yellow circular cursor (5 mm diameter) represented the participant’s hand position. Participants were instructed to guide the cursor into the start box and remain motionless. Once the cursor remained within the start box for at least 2 s with a velocity below 0.065 m/s, the start box disappeared and an auditory cue signaled the start of the trial. A white circular target (15 mm diameter) then appeared and moved along a pseudo-random sum-of-sines trajectory in both the *x*- and *y*-directions. The trajectory was generated by combining seven sinusoidal components along each axis (Fig. 1F). The target position *r*(*t*) at time *t* along each axis was defined as:

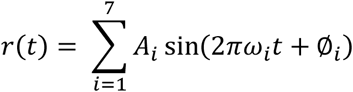

where *A*_*i*_, *ω*_*i*_ and ∅_*i*_ denote the amplitude, frequency, and phase of each sinusoidal component, respectively. Distinct amplitudes and frequencies were assigned to the *x*- and *y*-axes to dissociate hand responses to target motion along the two axes. The parameters were:

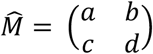

Frequencies were selected as prime multiples of 0.05 Hz to avoid harmonic overlap. Amplitudes were adjusted to maintain approximately constant peak velocity across frequencies while keeping movements within a comfortable tracking range (Yang et al., 2021; Zimmet et al., 2020). This design produced a temporally complex trajectory that repeated every 20 s. The phase of each sinusoidal component was randomized between trials to generate a unique trajectory and minimize memorization of movement patterns. In the baseline and post-learning blocks, however, phases were held constant to allow comparison of after-effects across sessions.

Participants were instructed to maintain the cursor at the center of the target throughout the trial. Successful tracking was indicated by a change in target color to green, providing positive visual feedback intended to encourage hand movements at the same constituent frequencies of the target trajectory.

#### MA segment and cursor jump

In all tracking trials, at a randomly selected time point (≥ 5 s after tracking initiation), the target temporarily ceased its sinusoidal motion and moved toward the lower-left corner of the workspace at a velocity matched to the mean velocity of its preceding motion. Upon reaching the corner, the target transitioned to a linear trajectory aligned with the MA and moved at either 30% or 70% of its mean velocity.

In approximately half of the trials, an unexpected instantaneous 3 cm cursor jump was introduced during this MA segment to probe rapid online responses and assess feedback control (Kasuga et al., 2015; Telgen et al., 2014). The jump occurred after the target had traveled 3–7 cm along the MA and was applied orthogonally, with half of the jumps directed +90° and half −90° relative to the MA. This design allowed participants to reach a steady movement state prior to the jump while reducing predictability of jump timing and direction.

Following the MA segment, the target resumed its original sinusoidal trajectory until the end of the trial. The cursor then returned linearly to its pre-jump position relative to the hand. The MA segment extended the 40 s sinusoidal tracking period by an additional 3–12 s, resulting in total trial durations of approximately 43–52 s.

### Point-to-point movement trial

Each trial began with a white square start box (10 mm) presented at the center of the workspace (Fig. 1E). A yellow circular cursor (5 mm diameter) represented the participant’s hand position. Participants were instructed to guide the cursor into the start box and remain motionless. Once the cursor remained within the start box for 2 s with a velocity below 0.065 m/s, a white circular target (15 mm diameter) appeared at one of seven locations on a virtual ring (10 cm radius). Target angles were randomly selected from 0°, ±30°, ±60°, and ±90° relative to the MA, with 0° aligned to the MA.

Participants were instructed to perform a rapid reaching movement toward the target following an auditory Go cue delivered at target onset. Movements initiated more than 500 ms after the cue triggered an unpleasant auditory tone to encourage rapid responses. Upon leaving the start box, the cursor disappeared and remained hidden until movement completion to minimize online visual corrections. When the cursor crossed the virtual target ring, it reappeared to indicate movement termination. The cursor endpoint and target remained visible for 2 s, with successful target acquisition indicated by the target turning green.

Movements with a peak velocity below 0.39 m/s triggered an unpleasant auditory tone to encourage faster movements in subsequent trials. At the end of each trial, the target and cursor disappeared, leaving only the start box and a white ring indicating the hand’s distance from the start position. The ring radius decreased as the hand approached the start box, and once the hand entered the box, the cursor reappeared to initiate the next trial.

### Analyses

All behavioral analyses were performed using custom MATLAB scripts (The MathWorks Inc., Natick, MA, USA).

#### Alignment analysis

For each tracking trial, hand and target trajectories were extracted and analyzed to quantify tracking performance. To assess the relationship between target motion and hand motion, a 2 × 2 alignment matrix was computed that optimally transformed the target trajectory *T* = (*T*_*x*_, *T*_*y*_) into the hand trajectory *H* = (*H*_*x*_, *H*_*y*_) (Yang et al., 2021). The alignment matrix was parameterized as:

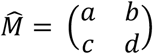

The matrix elements were estimated by minimizing the mean squared error (MSE) between the transformed target trajectory and the observed hand trajectory according to:

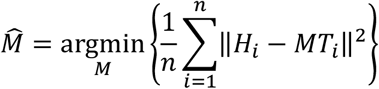

where *n* denotes the number of trajectory samples. Optimization was initialized with the identity matrix. The resulting alignment matrix 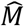 quantified how participants transformed their hand movements to track the target under the MR perturbation.

To quantify learning along the axis orthogonal to the MA, an orthogonal scaling factor (*S*) was computed from the alignment matrix. A unit vector orthogonal to the MA 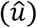 was transformed by the alignment matrix, and the orthogonal scaling factor was defined as:

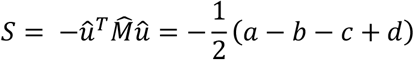

This orthogonal scaling factor ranged from −1 to 1, where −1 reflected baseline performance (no learning) and 1 indicated ideal learning of the MR mapping (perfect reversal).

#### Frequency analysis

For each tracking trial, the hand and target trajectories were interpolated and resampled at the trial’s mean sampling rate to obtain uniformly spaced samples. DC offsets were removed, and the discrete Fourier transform (DFT) of each signal was computed using MATLAB’s fast Fourier transform (FFT) algorithm. This yielded Fourier transforms of the hand *H*(*ω*) = (*H*_*x*_(*ω*), *H*_*y*_(*ω*)) and target *T*(*ω*) = (*T*_*x*_(*ω*), *T*_*y*_(*ω*)), represented as complex-valued phasors containing amplitude and phase information at each frequency. To characterize the frequency-dependent relationship between hand and target motion, frequency-specific transfer functions were computed by taking the ratio between hand and target phasors at the target frequencies. For *x*-target frequencies, the ratios *H*_*x*_(*ω*)/*T*_*x*_(*ω*) and *H*_*y*_(*ω*)/*T*_*x*_(*ω*) were calculated, whereas for *y*-target frequencies, the ratios *H*_*x*_(*ω*)/*T*_*y*_(*ω*) and *H*_*y*_(*ω*)/*T*_*y*_ (*ω*) were calculated. Gain was defined as the magnitude of each ratio, and phase difference was defined as its angle. Assuming similar behavior across adjacent *x*- and *y*-axis frequencies, adjacent frequency pairs were grouped to construct seven 2 × 2 gain matrices *G* spanning low to high frequencies: [(0.1, 0.15), (0.25, 0.35), (0.55, 0.65), (0.85, 0.95), (1.15, 1.45), (1.55, 1.85), (2.05, 2.15)]. For each frequency pair (*ω*_*x*_, *ω*_*y*_), the gain matrix *G*_*i*_ was defined as:

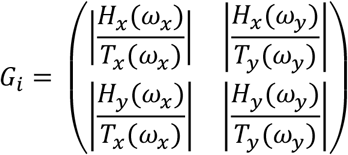

From each gain matrix, an orthogonal gain *g*_*i*_ was computed as:

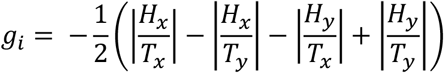

where all terms were evaluated at the corresponding frequency pair. Orthogonal gain values around −1 reflected baseline-like tracking behavior (no learning), whereas values around 1 indicated ideal learning of the MR mapping (perfect reversal).

#### Calculation of after-effects and learning level

After-effects and learning levels were computed using the orthogonal scaling factor, the frequency-specific orthogonal gain, or the mean off-diagonal components of the alignment matrix, depending on the analysis context. After-effects were quantified as the change in performance between the first post-learning trial and baseline performance:

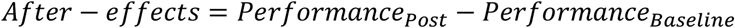

where *Performance*_*Post*_ denotes performance in the first post-learning trial (Day 1 or Day 5), and *Performance*_*Baseline*_ denotes the mean performance across all trials in the baseline block on Day 1. Late learning was quantified as the percentage improvement relative to baseline performance:

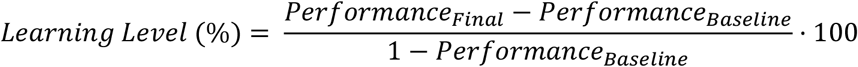

where *Performance*_*Final*_ denotes the mean performance during the final learning block on Day 5.

#### Hand response to cursor jump analysis

To assess how participants adjusted their corrective responses to unexpected cursor jumps, hand trajectories were analyzed within a time window spanning from jump onset to 1200 ms afterward. Trajectories were averaged within each experimental block to obtain the corrective hand response (*h*). Corrective responses were compared to an ideal corrective response (*h*_*ideal*_), representing the response expected after complete learning of the MR mapping. For each participant, *h*_*ideal*_ was defined as the mirrored version of their mean baseline corrective response from Day 1. A hand gain factor (*S*_*h*_) was then computed to quantify the alignment between the observed and ideal corrective responses:

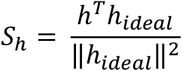

A value of −1 indicated alignment with the baseline corrective response, whereas a value of 1 indicated full alignment with the ideal corrective response under the MR mapping.

#### Point-to-point reaching analysis

For each trial, movement onset was defined as the first time point at which tangential velocity exceeded 5% of the trial’s peak velocity. The initial movement direction was calculated from the mean hand position within the 100–150 ms interval following movement onset. Initial error was then calculated as the absolute angular deviation between the initial movement direction and target angle.

#### Statistical analyses

All statistical analyses were conducted using MATLAB (The MathWorks Inc., Natick, MA, USA) and GraphPad Prism 10.4.0 (GraphPad Software, Boston, MA, USA). Statistical significance was set at *α* = 0.05. Directional one-sample tests were used only when the direction of the effect was specified a priori. All other tests were two-tailed. Holm–Šídák correction was applied within each family of multiple comparisons. Effect sizes for repeated-measures ANOVAs are reported as partial eta squared 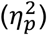, and effect sizes for one-sample tests of participant-level slopes are reported as Cohen’s *d*_*z*_. For repeated-measures ANOVAs with more than two levels, sphericity was assessed using Mauchly’s test. When sphericity was violated, Greenhouse–Geisser-corrected degrees of freedom and *p*-values were reported. Follow-up comparisons used the pooled residual error from the corresponding ANOVA.

Baseline tracking under veridical mapping was evaluated using a one-sample left-tailed *t*-test comparing the Day 1 baseline orthogonal scaling factor with zero. Changes in orthogonal scaling factor across practice were assessed using a one-way repeated-measures ANOVA with practice day (Days 1–5) as the within-subject factor. Holm–Šídák correction was applied across all 10 pairwise comparisons between practice days, although only adjacent-day comparisons were reported. Successful expression of the reversed mapping at the end of practice was evaluated using a one-sample right-tailed *t*-test comparing the orthogonal scaling factor from the final learning block on Day 5 with zero. Persistence of the learned mapping after restoration of veridical feedback was assessed using a one-way repeated-measures ANOVA comparing the Day 1 baseline average with the first post-learning trials on Days 1 and 5. The same analysis was applied to the mean off-diagonal alignment measure, defined as the average of the two off-diagonal elements of the alignment matrix.

For frequency-domain analyses, baseline orthogonal gain was averaged across Day 1 baseline trials separately for each of the seven frequency components. Baseline gains were compared with zero using one-sample left-tailed *t*-tests, with Holm–Šídák correction across the seven frequencies. Orthogonal gains in the final learning block on Day 5 were compared with zero using one-sample right-tailed *t*-tests, again with correction across the seven frequencies. Differences in normalized learning level across frequencies were assessed using a one-way repeated-measures ANOVA with frequency as the within-subject factor. Frequency-specific after-effects were defined as the difference between orthogonal gain on the first post-learning trial and the corresponding Day 1 baseline gain. After-effects were tested against zero at each frequency using right-tailed one-sample *t*-tests. Holm–Šídák correction was applied separately across the seven Day 1 tests and across the seven Day 5 tests. Changes in after-effects across practice were assessed using a two-way repeated-measures ANOVA with time (Day 1 and Day 5) and frequency (f_1_–f_7_) as within-subject factors.

#### Linear regression analyses

Frequency-dependent trends were evaluated at the participant level for baseline orthogonal gain, end-of-practice learning level, and after-effects on Days 1 and 5. For each dependent measure, a linear model was fitted separately to each participant’s values across the seven frequency conditions. The resulting participant-specific slopes were tested against zero using two-tailed one-sample t-tests. Complementary descriptive regressions were fitted to the seven group-averaged values for baseline orthogonal gain, end-of-practice learning level, after-effects on Days 1 and 5, and the Day 5-minus-Day 1 change in after-effects. Regression slopes were assessed using the model F-test, and goodness of fit was summarized using *R*^2^.

#### Exponential models for learning

To characterize learning dynamics, single-phase exponential models were fitted to block-averaged data for each learning measure. Orthogonal gain and hand corrective responses (hand gain factor) were modeled using a one-phase rising exponential association function:

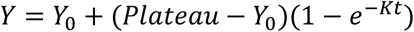

Initial angular error during point-to-point reaching, which decreased with learning, was modeled using a decaying exponential function:

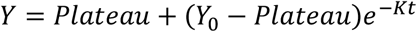

In both models, *Y*_0_ represents the initial value, *Plateau* denotes the asymptotic value approached with practice, *t* corresponds to time, and *K* defines the learning rate. The time constant *τ* = 1/*K* was used to compare learning dynamics across measures. Curve fitting was performed using least-squares regression without outlier exclusion, and goodness of fit was quantified using *R*^2^. Differences in time constants were evaluated using bootstrap resampling (3,000 resamples) to obtain 95% confidence intervals. After fitting, curves were normalized to their respective [*Y*_0_, *Plateau*] ranges to allow direct comparison across measures. For visualization in Figure 4E, the initial error and its corresponding decaying exponential fit were inverted and plotted as rising functions. Correspondence between learning measures was assessed by comparing their normalized values to the identity line (Figs. 4F and 4G, left panels). Deviation from perfect correspondence was quantified using RMSE, and paired *t*-tests were conducted to compare RMSE values between the low- and high-frequency conditions (f_1_ vs. f_7_).

### Computational model

We implemented a hybrid cortical-cerebellar neural network model controlling a two-joint planar arm. The model comprised interacting feedforward and feedback control pathways that jointly generated the motor command sent to the plant:

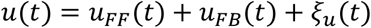

where *u*_FF_(*t*) ∈ ℝ^2^ denotes the predictive motor command generated by a cortical-like feedforward RNN, *u*_FB_(*t*) ∈ ℝ^2^ denotes the corrective motor command generated by a cerebellar-like feedback network, and *ξ*_*u*_(*t*) denotes zero-mean Gaussian motor noise. The feedforward RNN received target kinematics and generated predictive motor commands, whereas the feedback controller received delayed sensory error signals and generated online corrective commands.

Because sensory feedback was delayed, the controller maintained an internal estimate of the current arm state, *ŝ* (*t*), using a state estimator that integrated efference copy and delayed sensory observations. Mismatches between this estimate and the delayed sensory input generated sensory prediction errors that updated the estimator and drove the feedback network. The feedback pathway also provided a feedback-derived teaching signal used to update recurrent cortical weights of the feedforward RNN through a local Hebbian-style plasticity rule (Feulner et al., 2025).

#### Cortical RNN feedforward controller

The cortical feedforward controller was implemented as a continuous-time RNN. With recurrent state *x*(*t*) ∈ ℝ^*N*^and rectified linear-unit activation function *ϕ*(⋅), the network dynamics were defined as:

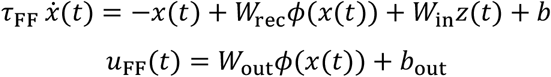

Inputs to the network included target position, target velocity, and task context:

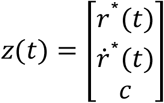

where *c* ∈ {0,1} indicated the current visuomotor mapping (0: veridical mapping; 1: MR mapping).

#### Cerebellar feedback controller

The cerebellar feedback controller was implemented as a neural network receiving visual and proprioceptive error signals. Visual and proprioceptive channels were modeled separately using distinct sensory delays (*d*_vis_ and *d*_prop_).

From the estimated state *ŝ* (*t*), estimated hand and cursor positions were computed as:

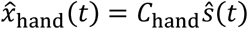

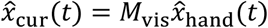

where *C*_hand_ maps state to hand position and *M*_vis_ denotes the visuomotor mapping (veridical or MR). Desired hand position under the current visuomotor mapping was defined as:

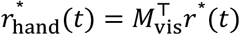

For the MR mapping,

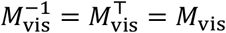

Visual and proprioceptive errors were computed as:

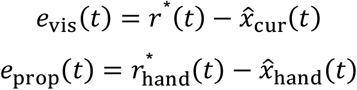

For each modality *k* ∈ {vis, prop}, low-pass filtered and derivative error signals were computed as:

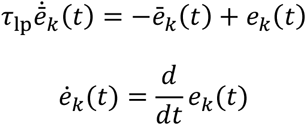

Feature vectors were constructed from instantaneous, filtered, and derivative error terms:

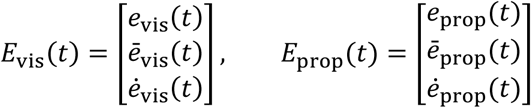

Visual and proprioceptive feature vectors were concatenated:

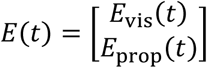

The combined feature vector was scaled using a diagonal gain matrix:

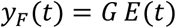

where *G* is a diagonal gain matrix (Table 1). Task context *c* was provided to the feedback-network input:

**Table 1.**
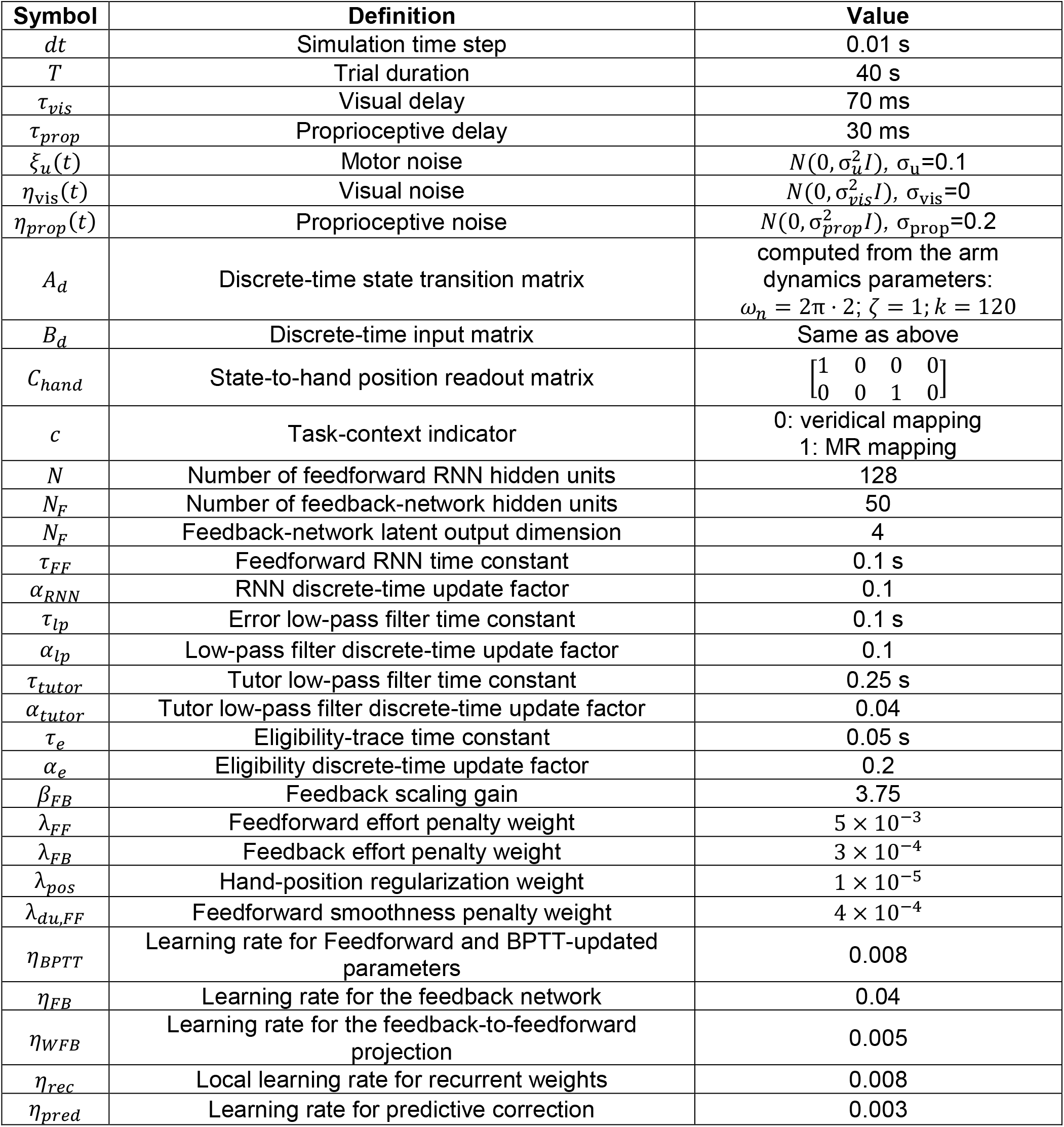
Model parameters.

| Symbol | Definition | Value |
| --- | --- | --- |
| $dt$ | Simulation time step | 0.01 s |
| $T$ | Trial duration | 40 s |
| $\tau_{vis}$ | Visual delay | 70 ms |
| $\tau_{prop}$ | Proprioceptive delay | 30 ms |
| $\xi_u(t)$ | Motor noise | $N(0, \sigma_u^2 I)$, $\sigma_u=0.1$ |
| $\eta_{vis}(t)$ | Visual noise | $N(0, \sigma_{vis}^2 I)$, $\sigma_{vis}=0$ |
| $\eta_{prop}(t)$ | Proprioceptive noise | $N(0, \sigma_{prop}^2 I)$, $\sigma_{prop}=0.2$ |
| $A_d$ | Discrete-time state transition matrix | computed from the arm dynamics parameters:<br>$\omega_n = 2\pi \cdot 2$; $\zeta = 1$; $k = 120$ |
| $B_d$ | Discrete-time input matrix | Same as above |
| $C_{hand}$ | State-to-hand position readout matrix | $\begin{bmatrix} 1 & 0 & 0 & 0 \\ 0 & 0 & 1 & 0 \end{bmatrix}$ |
| $c$ | Task-context indicator | 0: veridical mapping<br>1: MR mapping |
| $N$ | Number of feedforward RNN hidden units | 128 |
| $N_F$ | Number of feedback-network hidden units | 50 |
| $N_F$ | Feedback-network latent output dimension | 4 |
| $\tau_{FF}$ | Feedforward RNN time constant | 0.1 s |
| $\alpha_{RNN}$ | RNN discrete-time update factor | 0.1 |
| $\tau_{lp}$ | Error low-pass filter time constant | 0.1 s |
| $\alpha_{lp}$ | Low-pass filter discrete-time update factor | 0.1 |
| $\tau_{tutor}$ | Tutor low-pass filter time constant | 0.25 s |
| $\alpha_{tutor}$ | Tutor low-pass filter discrete-time update factor | 0.04 |
| $\tau_e$ | Eligibility-trace time constant | 0.05 s |
| $\alpha_e$ | Eligibility discrete-time update factor | 0.2 |
| $\beta_{FB}$ | Feedback scaling gain | 3.75 |
| $\lambda_{FF}$ | Feedforward effort penalty weight | $5 \times 10^{-3}$ |
| $\lambda_{FB}$ | Feedback effort penalty weight | $3 \times 10^{-4}$ |
| $\lambda_{pos}$ | Hand-position regularization weight | $1 \times 10^{-5}$ |
| $\lambda_{du,FF}$ | Feedforward smoothness penalty weight | $4 \times 10^{-4}$ |
| $\eta_{BPTT}$ | Learning rate for Feedforward and BPTT-updated parameters | 0.008 |
| $\eta_{FB}$ | Learning rate for the feedback network | 0.04 |
| $\eta_{WFB}$ | Learning rate for the feedback-to-feedforward projection | 0.005 |
| $\eta_{rec}$ | Local learning rate for recurrent weights | 0.008 |
| $\eta_{pred}$ | Learning rate for predictive correction | 0.003 |

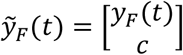

The feedback network was implemented as a one-hidden-layer neural network:

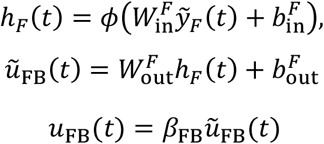

#### State estimator with predictive correction

The estimator predicted the latent arm state using an internal forward model of the plant dynamics:

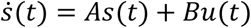

The estimator uses the same dynamics for state prediction:

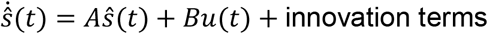

Visual and proprioceptive observations were delayed:

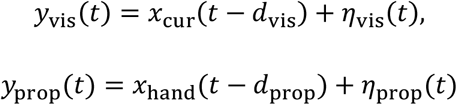

Sensory prediction errors were computed relative to delayed predicted states:

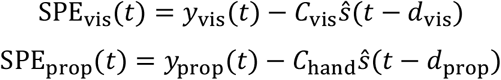

where *C*_vis_ = *M*_vis_*C*_hand_

The estimator incorporated these prediction errors using fixed observer gains:

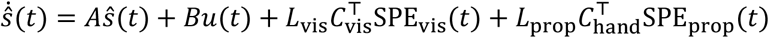

In addition to the fixed-gain observer, the model included a learned predictive correction term that compensated for systematic delay-induced mismatch between *y*_vis_(*t*) and the forward prediction. This nonlinear basis expansion of recent sensorimotor history was computed as:

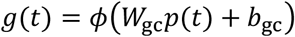

where *p*(*t*) concatenated recent motor commands (efference copy) across a delay window, recent visual cursor observations, and task context. The predictive module generated a state correction term:

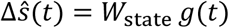

which was added to the delayed predicted state in the visual innovation term:

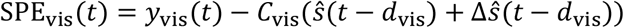

Correction weights were updated online from visual prediction error using a local delta rule:

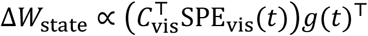

Learning of the predictive correction module was masked during cursor-jump epochs.

#### Arm dynamics and sensory delays

The arm was modeled as a stable second-order linear system in each dimension:

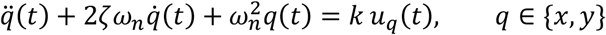

Equivalently,

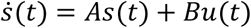

With hand position given by:

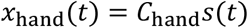

Cursor position was generated by applying the visuomotor transformation:

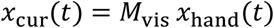

#### Local plasticity rule

Cortical recurrent weights were updated using a local learning rule combining an eligibility trace with a delayed teaching signal. The eligibility trace evolved according to:

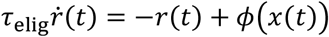

The learning signal was generated from feedback-network activity:

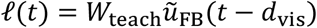

Recurrent weight updates were computed as:

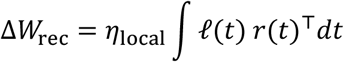

The feedback pathway additionally provided a tutor signal defined as a delay-aligned low-pass filtered version of the feedback command:

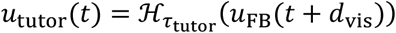

where ℋ_*τ*_ denotes a first-order low-pass filter. Tutor learning was masked during cursor-jump epochs.

#### Loss functions

Tracking loss was defined on the displayed cursor trajectory:

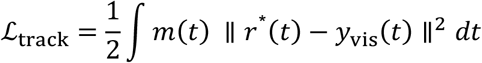

where *m*(*t*) excluded cursor-jump epochs.

Regularization terms penalized motor effort, feedforward smoothness, and workspace position:

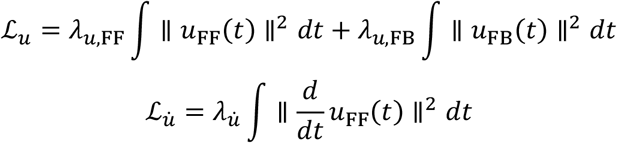

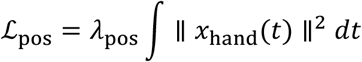

During MR learning, a tutor imitation loss was added:

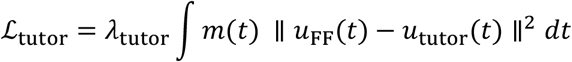

The total loss was:

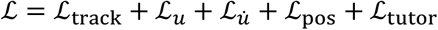

#### Task structure and training procedure

The target trajectory was modeled as a continuous two-dimensional sum-of-sines trajectory with seven frequencies per axis:

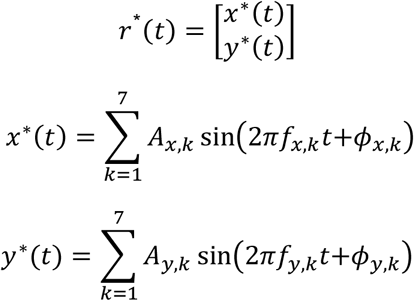

Frequency and amplitude parameters matched those used in the behavioral experiment. Phases were randomized independently for each trial. To enable controlled comparisons across simulations and ablation conditions, pseudo-random seeds used for trajectory generation were fixed across simulations. Consequently, the sequence of target trajectories was identical across simulations, whereas network initialization and motor noise were independently generated for each simulation.

MR was applied only to the displayed cursor mapping:

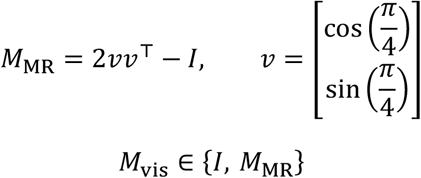

where *I* denotes the identity mapping and *M*_MR_ denotes reflection across the MA aligned at 45°.

Each simulation consisted of pretraining, baseline training, MR training, and post-learning phase, with a fixed visual delay present throughout. During pretraining, the model acquired stable tracking under the veridical mapping while experiencing delayed visual feedback. Baseline training continued under the veridical mapping to further reduce tracking error and stabilize performance. During MR practice, the visuomotor mapping was switched to *M*_MR_ while maintaining the same visual delay. During post-learning phase, the visuomotor mapping was restored to the veridical condition.

To quantify pathway-specific contributions, probe forward passes were interleaved during MR practice without weight updates. In feedback-only probe trials, the feedforward motor command was set to zero such that behavior was generated exclusively by the feedback pathway. In feedforward-only probe trials, the feedback motor command was set to zero. These probe conditions isolated pathway-specific motor output while preserving the same target trajectories and internal network dynamics used during the full model simulations.

Parameter values used in the simulations are summarized in Table 1.

#### Evaluation metrics of the model

##### Tracking accuracy

Tracking performance was quantified using the MSE between target and the cursor position:

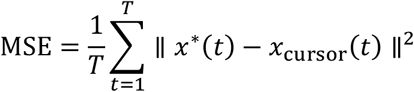

RMSE was additionally computed as:

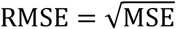

##### Frequency-dependent performance

Frequency-domain transfer matrices relating target and hand motion were computed separately for each target frequency component using the same frequency-domain analysis applied to the behavioral data (see Frequency analysis). Orthogonal gain was then computed from the resulting transfer matrices to quantify behavior at each frequency.

##### Frequency-specific after-effects

Orthogonal gain was computed separately for each target frequency f_k_ during baseline and post-learning phase. Baseline performance was defined as the mean orthogonal gain across the final 20 baseline trials 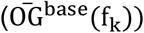. Post-learning performance was defined as orthogonal gain on the first post-learning trial immediately following removal the MR perturbation 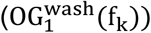. After-effect magnitude was computed as the baseline-corrected change following perturbation removal:

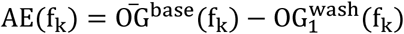

##### Alignment of recurrent plasticity with the teaching signal

To quantify the alignment between recurrent plasticity and the local Hebbian teaching update, cosine similarity was computed between the realized recurrent weight change and the local teaching-rule update on each trial during MR learning.

On trial *n*, the local teaching-rule update was computed as a trial-summed outer product between a feedback-derived postsynaptic signal and a presynaptic eligibility trace:

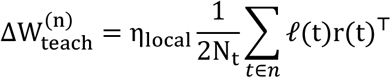

where *ℓ*(t) ∈ ℝ^N^ denotes the feedback-derived postsynaptic teaching signal, r(t) ∈ ℝ^N^ denotes the presynaptic eligibility trace, and N_t_ denotes the number of time points included in the update.

The realized recurrent weight change during trial *n* was defined as:

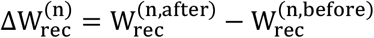

Cosine similarity between the vectorized matrices was computed as:

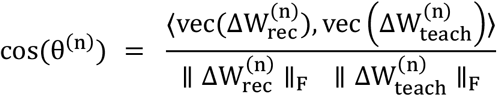

where ⟨⋅,⋅⟩ denotes the dot product and ∥⋅∥_F_ denotes the Frobenius norm. Cosine similarity was computed on each trial during MR learning and averaged across simulations.

Low-rank structure of recurrent plasticity:

To assess the dimensionality of recurrent plasticity, singular value decomposition (SVD) was applied to the trial-wise recurrent weight change matrix:

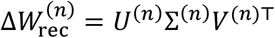

For each trial, trial-averaged postsynaptic teaching and presynaptic eligibility directions were computed as:

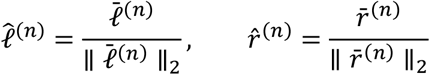

Alignment between the dominant singular modes and these directions was quantified using cosine projections:

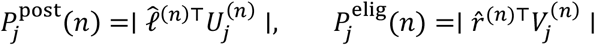

Here 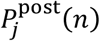 measures whether the left singular vectors ( 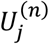; postsynaptic structure of the update) align with the teaching-signal direction, and 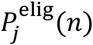 measures whether the right singular vectors ( 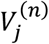; presynaptic structure of the update) align with the eligibility-trace direction. Analyses focused on the leading singular modes (*j* = 1, …,5), and **projections were averaged across simulations**.

## Notes

### Competing Interest Statement

The authors have declared no competing interest.

### Summary of Updates

New analysis, new statistical analysis, revised text and revised figures.

### Funding

- Israel Science Foundation: 2484/23
- United States-Israel Binational Science Foundation: 2021323

