## Supplementary Information for "Feedback–feedforward dynamics shape continuous motor learning"

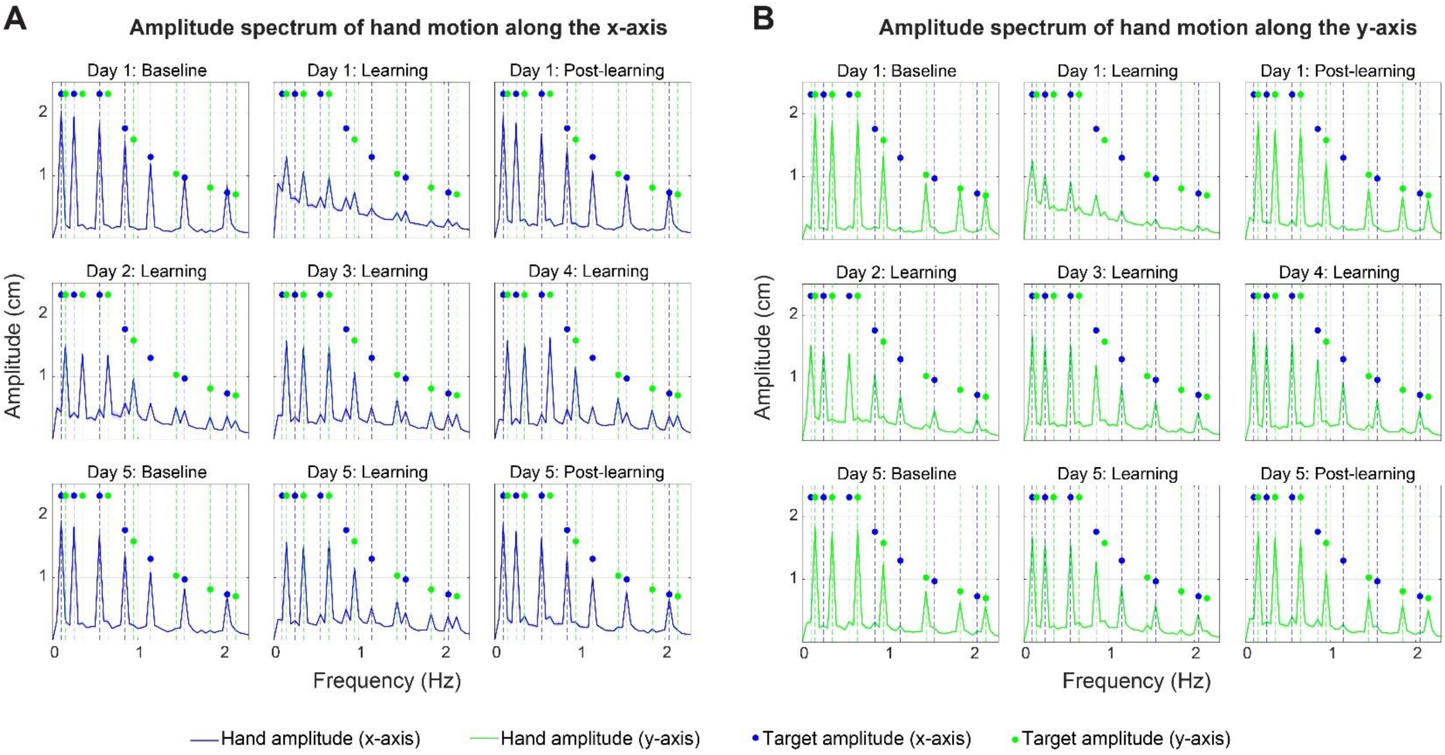


**Figure S1. Hand motion amplitude spectrum along the (A) $x$-axis and (B) $y$-axis.** Each subplot corresponds to a different experimental phase across five learning days. Solid lines represent averaged hand amplitude spectra across participants, with shaded areas indicating SEM. Blue and green circles mark target amplitudes at designated frequencies along the $x$- and $y$-axes, respectively. Dashed vertical lines indicate the target frequencies to facilitate visual interpretation.

Under veridical mapping, participants predominantly moved at the frequencies composing the target trajectory along each corresponding axis, consistent with the response of a linear system, in which the output comprises the same frequencies as the input. With practice under mirror reversal mapping, the amplitude spectrum gradually shifted, showing increased amplitudes at frequencies aligned with the orthogonal axis. For example, the hand amplitude spectrum along the $y$-axis (green line, Figure S2B) at baseline closely matched the target amplitude spectrum along the same axis (green dots), indicating axis-specific tracking. However, by Day 5 of learning under the mirror reversal mapping, there is a clear shift: the hand's amplitude spectrum along the $y$-axis increasingly aligns with the $x$-axis target frequencies (blue dots). This shift reflects the learning of the altered visuomotor mapping, where participants learned to produce hand movements along the orthogonal axis to achieve accurate cursor tracking. Despite this spatial remapping, participants' movements remained confined to the set of target frequencies along both axes, indicating that responses remained spectrally constrained to the input, a characteristic feature of a linear system.

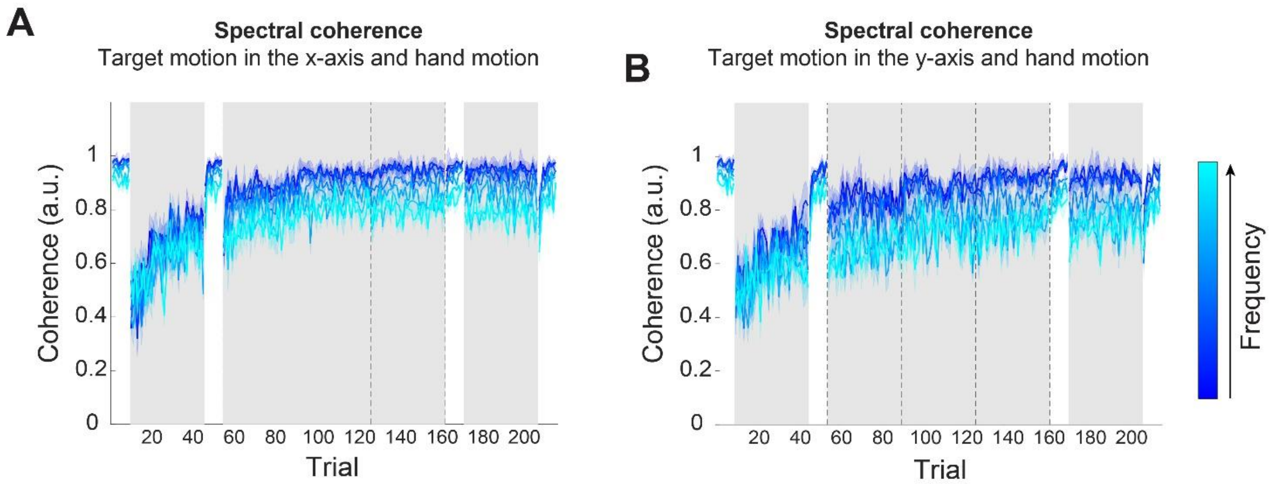


**Figure S2. Spectral coherence between target and hand motion. (A)** Coherence between target motion along the $x$-axis and hand motion in both axes. **(B)** Coherence between target motion along the $y$-axis and hand motion in both axes. Each line represents the coherence at a specific target frequency, averaged across participants. Shaded areas indicate the SEM. Gray background regions denote learning phase under mirror reversal mapping.

To quantify the linear relationship between target and hand motion, we computed spectral coherence on a trial-by-trial basis. For each participant, continuous trajectories of the target and hand were extracted from individual trials across five experimental days. Coherence was computed separately for target motion along the $x$- and $y$-axes, using hand motion in both axes as joint outputs in a single-input, multi-output framework. Coherence values were then interpolated at seven predefined target frequencies per axis to assess system behavior at task-relevant frequencies. During the baseline phase, coherence values were consistently high across all frequencies, indicating a strong linear relationship between target motion and hand responses (mean ± SEM: $x$-axis: $f_1$: 0.98 ± 0.002, $f_7$: 0.90 ± 0.01; $y$-axis: $f_1$: 0.97 ± 0.01, $f_7$: 0.89 ± 0.02). Introduction of the mirror reversal mapping on Day 1 led to a substantial decrease in coherence (mean ± SEM: $x$-axis: $f_1$: 0.67 ± 0.02, $f_7$: 0.60 ± 0.02; $y$-axis: $f_1$: 0.64 ± 0.03, $f_7$: 0.55 ± 0.03). However, coherence values were rapidly recovered with practice: by Day 2, coherence had increased (mean ± SEM: $x$-axis: $f_1$: 0.86 ± 0.03, $f_7$: 0.73 ± 0.03; $y$-axis: $f_1$: 0.84 ± 0.03, $f_7$: 0.65 ± 0.04), and approached baseline levels by Day 5 (mean ± SEM: $f_1$: $x$-axis: 0.95 ± 0.0049, $f_7$: 0.78 ± 0.02; $y$-axis: $f_1$: 0.92 ± 0.01, $f_7$: 0.72 ± 0.02). These results indicate that, while the mirror reversal transformation initially perturbed the linear relationship between target and hand motion, participants rapidly restored linear behavior across all frequencies through practice.

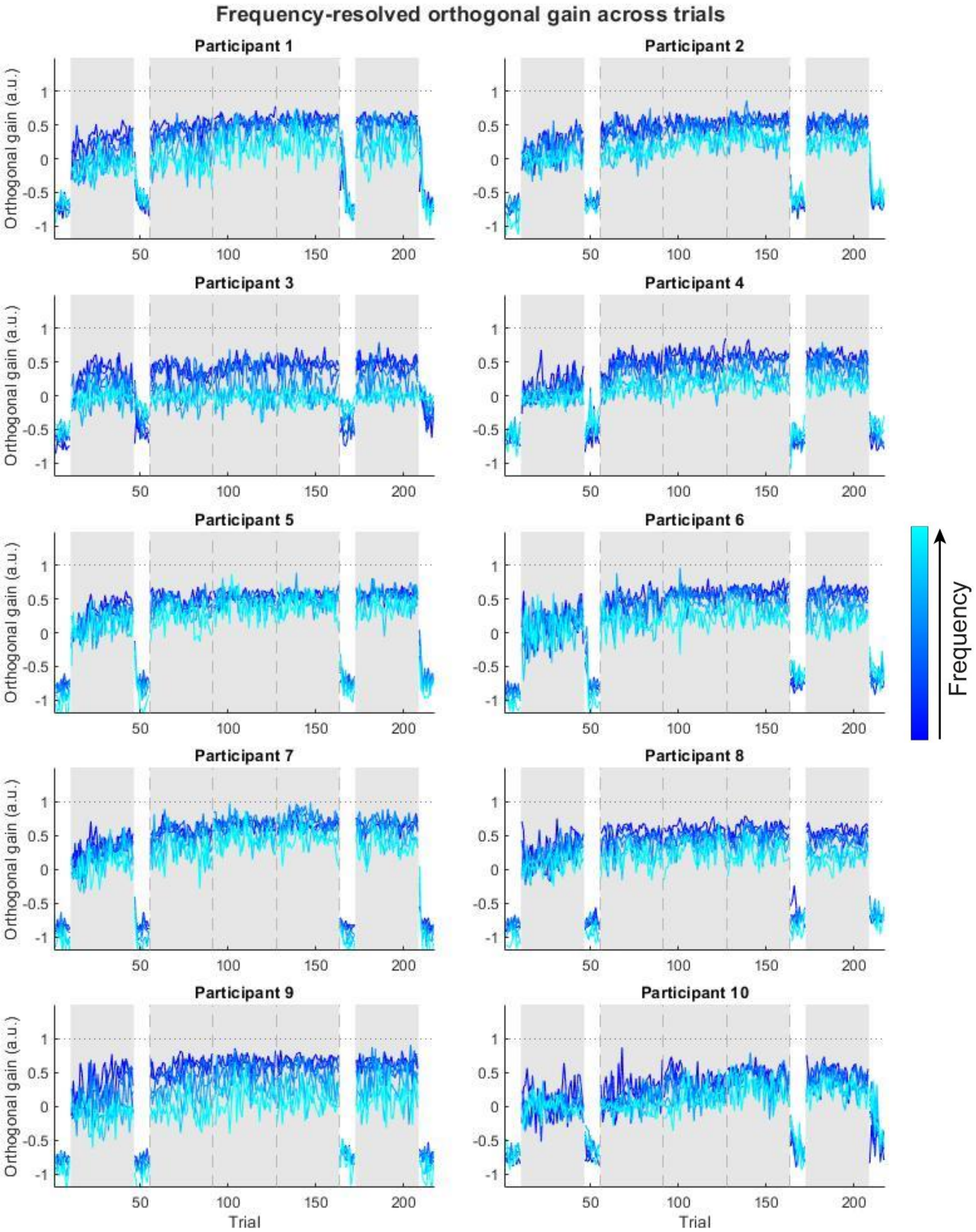


**Figure S3. Participant-level frequency-resolved orthogonal gain across practice.** Each panel shows orthogonal gain across experimental trials for one participant. Traces represent the seven frequency components ($f_1$–$f_7$). Gray background indicates practice under the mirror reversal mapping, and vertical dashed lines mark the boundaries between practice days. Traces are separated at transitions between baseline, learning, and post-learning phases.

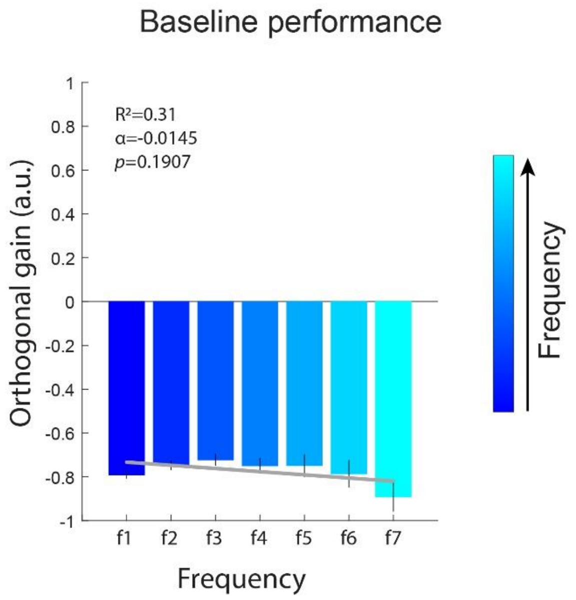


**Figure S4. Frequency-dependent orthogonal gain during baseline phase.** Orthogonal gain values were averaged across all trials in the baseline block on Day 1. Error bars represent the SEM. Regression analysis revealed no significant frequency-dependent variation, indicating that baseline performance was stable across frequencies.

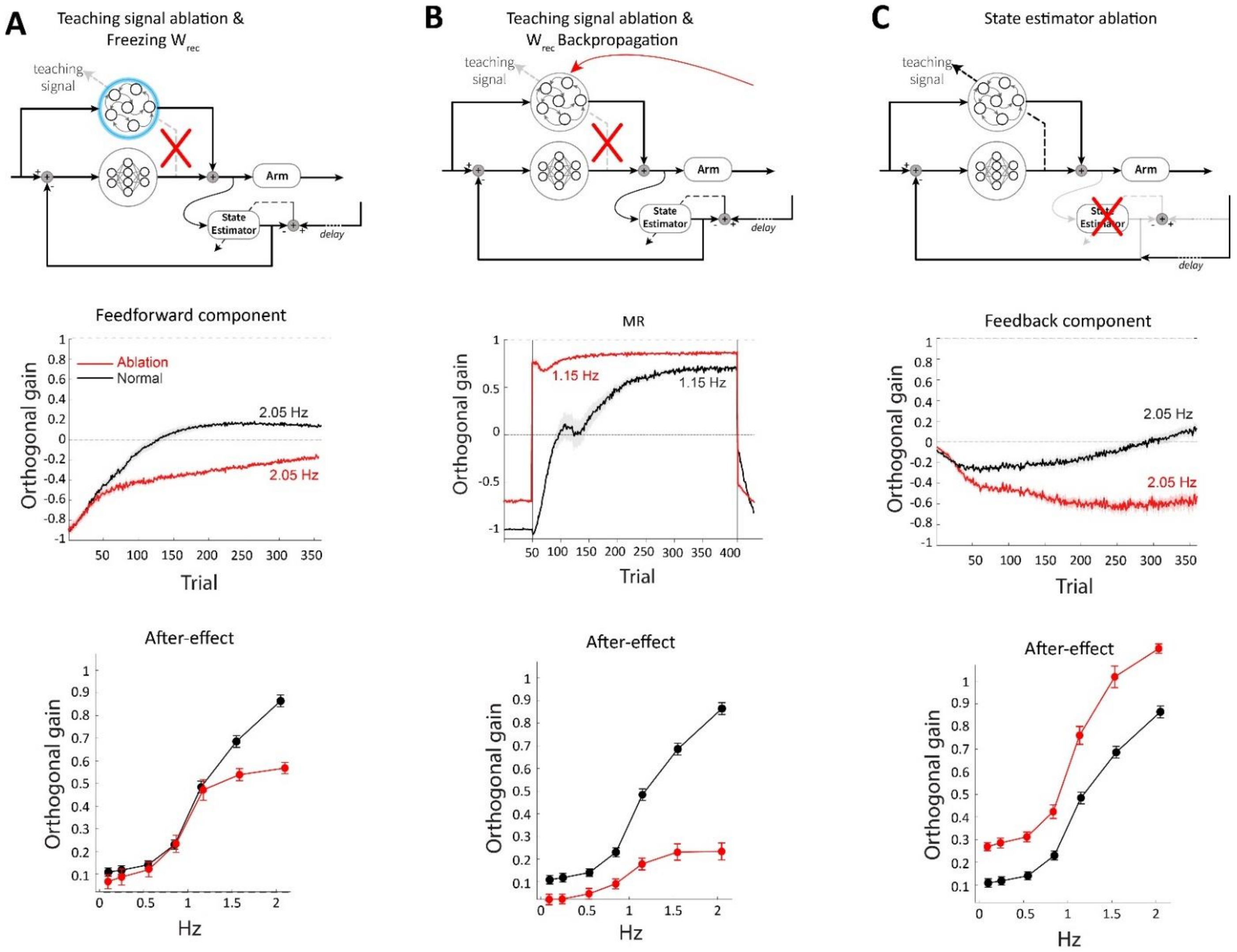


**Figure S5. Ablation analyses of the model identify distinct roles for teaching-driven recurrent plasticity and state estimation. (A)** Ablation of the feedback-derived teaching pathway together with freezing of cortical recurrent weights ($W_{rec}$) (top panel). Middle, feedforward component of orthogonal gain during mirror-reversal learning at a representative high frequency (2.05 Hz), shown for the intact model (black) and the ablated model (red). Bottom, after-effect as a function of frequency. Removing the teaching pathway and preventing recurrent adaptation strongly attenuated the emergence of the feedforward component at high frequency and reduced the frequency-dependent after-effect, indicating that feedback-derived tutoring and recurrent plasticity are required for the gradual buildup of predictive control. **(B)** Ablation of the local teaching pathway while allowing recurrent weights to be updated by backpropagation (top panel). Middle, total orthogonal gain during mirror-reversal learning at a representative intermediate frequency (1.15 Hz), for the intact model (black) and the ablated/backpropagation model (red). Bottom, after-effect as a function of frequency. Although backpropagation improved online task performance, it did not reproduce the characteristic temporal profile or persistence of learning expressed by the intact model, showing that generic optimization alone is insufficient to capture the observed dynamics. **(C)** Ablation of the state estimator (top panel). Middle, feedback component of orthogonal gain during mirror-reversal learning at a representative high frequency (2.05 Hz), for the intact model (black) and the ablated model (red). Bottom, after-effect as a function of frequency. Removing the state estimator drove the high-frequency feedback component in the wrong direction during learning and produced an exaggerated high-frequency after-effect, consistent with delay-dominated feedback corrections in the absence of predictive state estimation. Red traces indicate the ablation condition; black traces indicate the intact model. Error bars denote SEM across simulated networks (n=10).

To determine which components of the model are necessary to reproduce the behavioral signature observed in humans, we carried out three targeted ablation analyses. In the *first*, we removed the feedback-derived teaching pathway and froze recurrent cortical weights. Under this manipulation, the model retained limited online compensation, but the gradual emergence of the feedforward component was markedly reduced, and the after-effect remained substantially smaller, particularly at higher frequencies. This indicates that feedback-derived teaching signals and recurrent plasticity are both important for building the predictive component that supports later learning.

In the *second* analysis, we removed the local tutor-like recurrent learning mechanism but allowed recurrent weights to be optimized by backpropagation. This manipulation improved task performance during mirror-reversal training, showing that the task can still be solved in an algorithmic sense. However, the resulting trajectory no longer reproduced the gradual frequency-dependent progression or the persistence profile characteristic of the intact model. Thus, matching the behavioral signature requires more than successful optimization; it depends on the specific learning architecture through which feedback shapes feedforward control.

In the *third* analysis, we ablated the state estimator. This produced a qualitatively distinct pattern: at high frequency, the feedback component became strongly negative during mirror-reversal learning, consistent with delayed sensory errors driving corrections with an effectively inappropriate sign when not supported by prediction. At the same time, the after-effect became abnormally large at higher frequencies. Together, these results suggest that feedback serves as a tutor for the development of long-term cortical mappings, with the state estimator supporting this process by reducing delay-induced distortions in the feedback signal.
